# Cerebellar circuits for disinhibition and synchronous inhibition

**DOI:** 10.1101/2023.09.15.557934

**Authors:** Elizabeth P. Lackey, Luis Moreira, Aliya Norton, Marie E. Hemelt, Tomas Osorno, Tri M. Nguyen, Evan Z. Macosko, Wei-Chung Allen Lee, Court A. Hull, Wade G. Regehr

**Affiliations:** Department of Neurobiology, Harvard Medical School, Boston MA, United States; Department of Neurobiology, Duke University Medical School, Durham, United States; Broad Institute of Harvard and MIT, Stanley Center for Psychiatric Research, Cambridge, MA, USA; Kirby Neurobiology Center, Boston Children’s Hospital, Harvard Medical School, Boston, MA, USA

## Abstract

The cerebellar cortex contributes to diverse behaviors by transforming mossy fiber inputs into predictions in the form of Purkinje cell (PC) outputs, and then refining those predictions^1^. Molecular layer interneurons (MLIs) account for approximately 80% of the inhibitory interneurons in the cerebellar cortex^2^, and are vital to cerebellar processing^1,3^. MLIs are thought to primarily inhibit PCs and suppress the plasticity of excitatory synapses onto PCs. MLIs also inhibit, and are electrically coupled to, other MLIs^4–7^, but the functional significance of these connections is not known^1,3^. Behavioral studies suggest that cerebellar-dependent learning is gated by disinhibition of PCs, but the source of such disinhibition has not been identified^8^. Here we find that two recently recognized MLI subtypes^2^, MLI1 and MLI2, have highly specialized connectivity that allows them to serve very different functional roles. MLI1s primarily inhibit PCs, are electrically coupled to each other, fire synchronously with other MLI1s on the millisecond time scale *in vivo*, and synchronously pause PC firing. MLI2s are not electrically coupled, they primarily inhibit MLI1s and disinhibit PCs, and are well suited to gating cerebellar-dependent learning^8^. These findings require a major reevaluation of processing within the cerebellum in which disinhibition, a powerful circuit motif present in the cerebral cortex and elsewhere^9–17^, greatly increases the computational power and flexibility of the cerebellum. They also suggest that millisecond time scale synchronous firing of electrically-coupled MLI1s helps regulate the output of the cerebellar cortex by synchronously pausing PC firing, which has been shown to evoke precisely-timed firing in PC targets^18^.

---

Molecular layer interneurons (MLIs) play vital roles in cerebellar processing^1,3^. When mossy fibers convey signals from the rest of the brain and spinal cord, they activate granule cells that excite PCs, and excite MLIs that in turn disynaptically inhibit PCs. MLIs control calcium signaling in PC dendrites^19,20^, prevent the induction of long-term plasticity at granule cell to PC synapses^20^, and decrease PC firing^21^. Suppressing MLI firing degrades coordinated movement, suppresses learned motor responses^22^, impairs cerebellar-dependent motor learning^20,23–26^, and impairs reversal learning, novelty seeking and social behaviors^25^.

MLIs can contribute to cerebellar processing in many ways. All MLIs make conventional GABAergic synapses, but those located near the PC layer also make specialized structures known as pinceaux near the initial segments of PC axons to provide ephaptic inhibition^27^. MLIs are electrically coupled to each other^4–6^, leading to synchronous MLI firing on the millisecond time scale in brain slice^5,6,18^, but it is not known if MLIs fire synchronously in behaving animals. In addition to inhibiting PCs, MLIs inhibit other MLIs, but the role of such inhibition is not known. One intriguing possibility is that MLI-MLI inhibition could implement disinhibition, a powerful circuit used for computations such as selective gating and gain modulation elsewhere in the brain^9–14^. However, such disinhibition requires a specialized subpopulation of neurons that primarily inhibit other interneurons, and it is not known if such an MLI subpopulation exists. Using snRNA-seq, we recently found that MLIs are comprised of two molecularly distinct subtypes, which we named MLI1 and MLI2, that do not correspond to the classic basket cell and stellate cell categories^2^. There are approximately three times as many MLI1s as MLI2s intermingled throughout the molecular layer^2^, with a higher density of MLI1s near the PC layer^28^. MLI1 and MLI2 showed intriguing differences, such as in their excitability and expression of *Gjd2* (the gene encoding connexin 36), increasing the computational potential of MLIs, and suggesting that they could contribute to cerebellar processing in unexpected ways.

Here we determine the synaptic connectivity and electrical coupling of MLI1s and MLI2s, how they influence PC firing, and how they fire *in vivo*. We find that MLI1s are electrically coupled to each other, and they primarily inhibit PCs. In contrast, MLI2s are not electrically coupled, and they primarily inhibit MLI1s to disinhibit PCs. *In vivo* recordings suggest that MLI1s fire synchronously and provide precisely timed inhibition to suppress PC firing, whereas MLI2s promote PC firing. We conclude that specialized firing properties, electrical coupling and synaptic targeting allow MLI1 and MLI2 to have opposing influences on the PC outputs of the cerebellar cortex. This greatly expands the computational power of MLIs in cerebellar processing.

## Targeting MLI1 and MLI2 subtypes

It was necessary to discriminate between MLI1 and MLI2 subtypes in order to compare their synaptic connectivity. We explored the possibility of using transgenic mice to label subtypes of MLIs, based on the selective expression of *Nxph1* in MLI2s and *Gjd2* in MLI1s^2^. We made an *Nxph1^Cre^* mouse line to help identify MLI2s (**Extended Data Fig. 1a-c**), and used *Gjd2-EGFP* mice^29^ to help identify MLI1s (**Extended Data Fig. 1de**). Fluorophore expression was not restricted to a subtype for either of these mice, but in *Nxph1^Cre^ Ai14* mice and in *Gjd2-EGFP* mice the brightest cells were, respectively, MLI2 and MLI1.

*Nxph1^Cre^ Ai14* were particularly useful for targeting MLI2s. Ultimately, we relied on electrophysiological properties to identify MLI subtypes (**Extended Data Fig. 2**). We restricted our recordings to the inner two-thirds of the molecular layer where their electrical properties are most distinct, and we only determined the synaptic connectivity of MLIs that were unambiguously classified. Based on the finding that only MLI1s express *Gjd2* (connexin 36) and have spikelets (the characteristic currents arising from activity in electrically-coupled MLIs), we used spikelets to identify MLI1s, and a lack of spikelets combined with a high input resistance to identify MLI2s (**Extended Data Fig. 2a-c**). We found that for MLI1s and MLI2s identified by these criteria, MLI1s had lower input resistances, smaller I_h_, and that their membrane potential decayed more rapidly following an evoked action potential compared to MLI2s (**Extended Data Fig. 2c-h**). We also used a whole-cell recording pipette to fluorescently label MLIs that had been identified as either MLI1 or MLI2 (**Fig. 1**, **Extended Data Fig. 2ij**, **Fig. 3a**), and found that they had characteristic morphologies. Fluorescence images of MLI1-PC pairs show an MLI1 near the PC layer that looks like a classic basket cell with prominent collaterals that contribute to pinceaux (**Fig. 1a**), and an MLI1 further away from the PC layer that looks more like a classic stellate cell, except that it also extends two collaterals to the vicinity of the initial segments of PC axons to contribute to pinceaux (**Fig. 1b**). MLI2s had classic stellate cell morphologies, and lacked collaterals that make pinceaux-like structures below the PC layer (**Fig. 1gh**). For MLIs subtyped on the basis of their electrophysiological properties, we found that 10/10 MLI1s and 0/6 MLI2s had collaterals that contributed to pinceaux.

**Figure 1.**
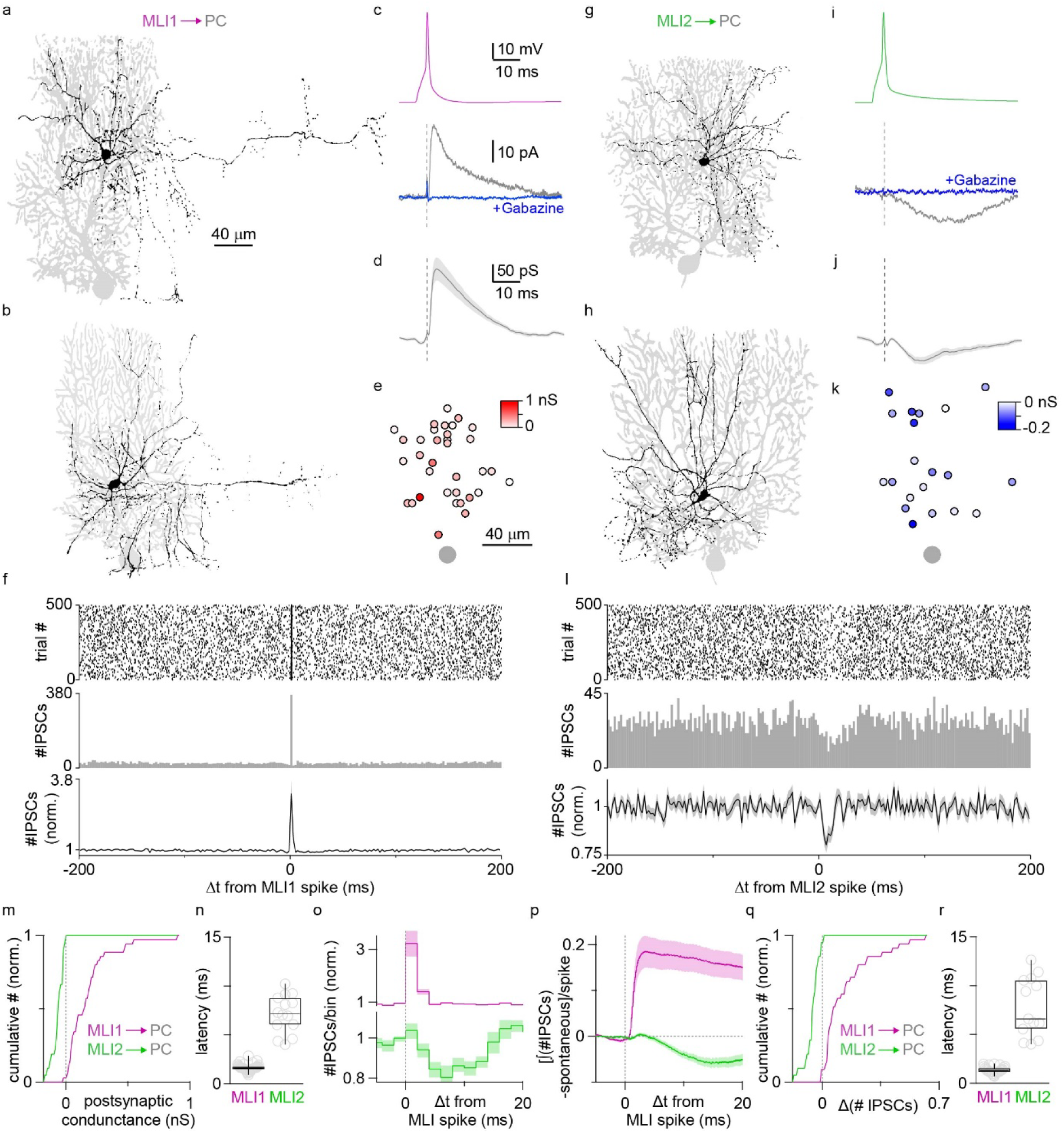
MLI1s powerfully inhibit Purkinje cells but MLI2s do not. Paired recordings between MLIs and PCs were performed, with the presynaptic MLI in current clamp and the synaptic responses in PCs measured in voltage clamp. **a.** Fluorescence image of an MLI1 located in the middle of the molecular layer (black) and PC (grey) pair. Scale bar also applies to **bgh**. **b.** Same as **a** but for an MLI1 located in the lower molecular layer. **c.** Paired recording from an MLI1 (purple) to PC (grey) pair. The GABA_A_R antagonist gabazine eliminated the synaptic response (*blue*). Scale bars also apply to **i**. Traces are averages of 500 trials, as is the case for all synaptic currents in Fig. 1 and Fig. 2. **d.** Average MLI1-PC synaptic currents. Scale bars also apply to **j**. **e.** Position of MLI1s relative to PCs color-coded for IPSC strength for all pairs. **f.** Raster plot of IPSCs (*top*) and corresponding histogram of IPSCs (*middle*) for an MLI1-PC cell pair, and average of histograms (*bottom*) for all MLI1-PC cell pairs. **g-l.** Same as **a-f** but for MLI2s. **m.** Cumulative plot of amplitudes of all MLI1-PC (*purple*, n=35, p=1E-10) and MLI2-PC (*green*, n=21, p=3E-09) synaptic responses. **n**. Latencies of synaptic responses. **o** Expanded histograms from **f** (*upper*) and **l** (*lower*). **p.** Average change in the integrated number of IPSCs per spike for all MLI-PC pairs. **q.** Cumulative plot summarizing the integrated number of IPSCs per stimulus for all pairs (p=9E-11). **r.** Latencies of change in integrated number of IPSCs per spike. See Fig. 3S for details on analysis.

## MLI1s inhibit PCs and MLI2s disinhibit PCs

Paired recordings indicate that MLI1s and MLI2s have very different effects on PCs. An inhibitory postsynaptic current (IPSC) is shown for an MLI1-PC pair (**Fig. 1c**), and the average conductance was 187 ± 31 pS for MLI1-PC pairs (**Fig. 1d**). The strengths of MLI1-PC synapses as a function of relative MLI1-PC positions are shown (**Fig. 1e**). MLI1s inhibited 94% of nearby PCs (**Extended Data Fig. 3a-c**, 33 of 35), with short latencies (**Fig. 1n**, 1.65 ± 0.07 ms). Spontaneous IPSCs were present at high frequencies in PCs (**Extended Data Fig. 3fg**, **Fig. 1f**), and MLI1 stimulation evoked large, brief increases in IPSC frequencies in PCs (**Extended Data Fig. 3h**, **Fig. 1fo**). MLI2s had very different effects on PCs. Rather than evoking an inhibitory outward current, MLI2s evoked an average inward current (excitatory) that was blocked by the GABA_A_R antagonist gabazine (**Fig. 1i-k**). Average inward currents were evoked in 71% of MLI2-PC pairs (**Extended Data Fig. 3de**, 15 of 21). Their long latencies (**Fig. 1n**) raised the possibility that MLI2s might suppress spontaneous inhibitory inputs onto PCs. This was confirmed by the observation that evoking a single spike in an MLI2 transiently reduced IPSC frequency in PCs (**Fig. 1lo**). In order to assess the effects of a single MLI spike on IPSCs in a PC, we detected IPSCs, integrated the events, and subtracted the spontaneous events, leaving behind the influence of stimulation (**Extended Data Fig. 3hi**, **Fig. 1p**). For all MLI1-PC pairs, MLI1s evoked a short-latency IPSC in approximately 18% of the trials (although this is an underestimate because we did not detect all events), and increases in IPSC frequency were evoked in 80% of MLI1-PC pairs (**Extended Data Fig. 3kl**, **Fig. 1o-q**, 28 of 35). For 71% of MLI2-PC pairs, IPSCs were suppressed with a long latency (7.82 ± 0.84 ms, n=15) (**Extended Data Fig. 3km**, **Fig. 1o, r**). These findings indicate that MLI1s inhibit most nearby PCs, and MLI2s disinhibit many nearby PCs.

## MLI2s powerfully inhibit MLI1s

The above observations suggested that MLI2s disinhibit PCs by inhibiting MLI1s. We therefore recorded from pairs of identified MLIs to assess the target dependence of synaptic connections, and we determined the electrical coupling between MLI subtypes. MLI2-MLI1 IPSCs were large and short latency (**Fig. 2a**). MLI2s inhibited a large percentage of nearby MLI1s (80%, 16/20, **Fig. 2j**), and the average peak inhibitory conductance for all cell pairs was 247 ± 63 pS (**Fig. 2b**). MLI1-MLI2 connections were present in 60% (12/20) of the pairs, (**Fig. 2cj**), but they were weak (22.1 ± 7.3 pS, **Fig. 2d**). Weak inhibition was also present in 50% (10/20) of MLI2-MLI2 connections (27.3 ± 9.1 pS, **Fig. 2efj**). Electrical coupling was present in 67% (20/30) of MLI1-MLI1 pairs, but was not observed for any MLI2-MLI1, MLI1-MLI2, and MLI2-MLI2 pairs (**Fig. 2gk**). For electrically-coupled MLI1s, the presynaptic action potential produced a large inward current in target cells that could be isolated by blocking GABA_A_ receptors with gabazine. Synaptic currents were quantified using the gabazine-sensitive component (**Fig. 2g**, *lower*, *black*).

**Figure 2.**
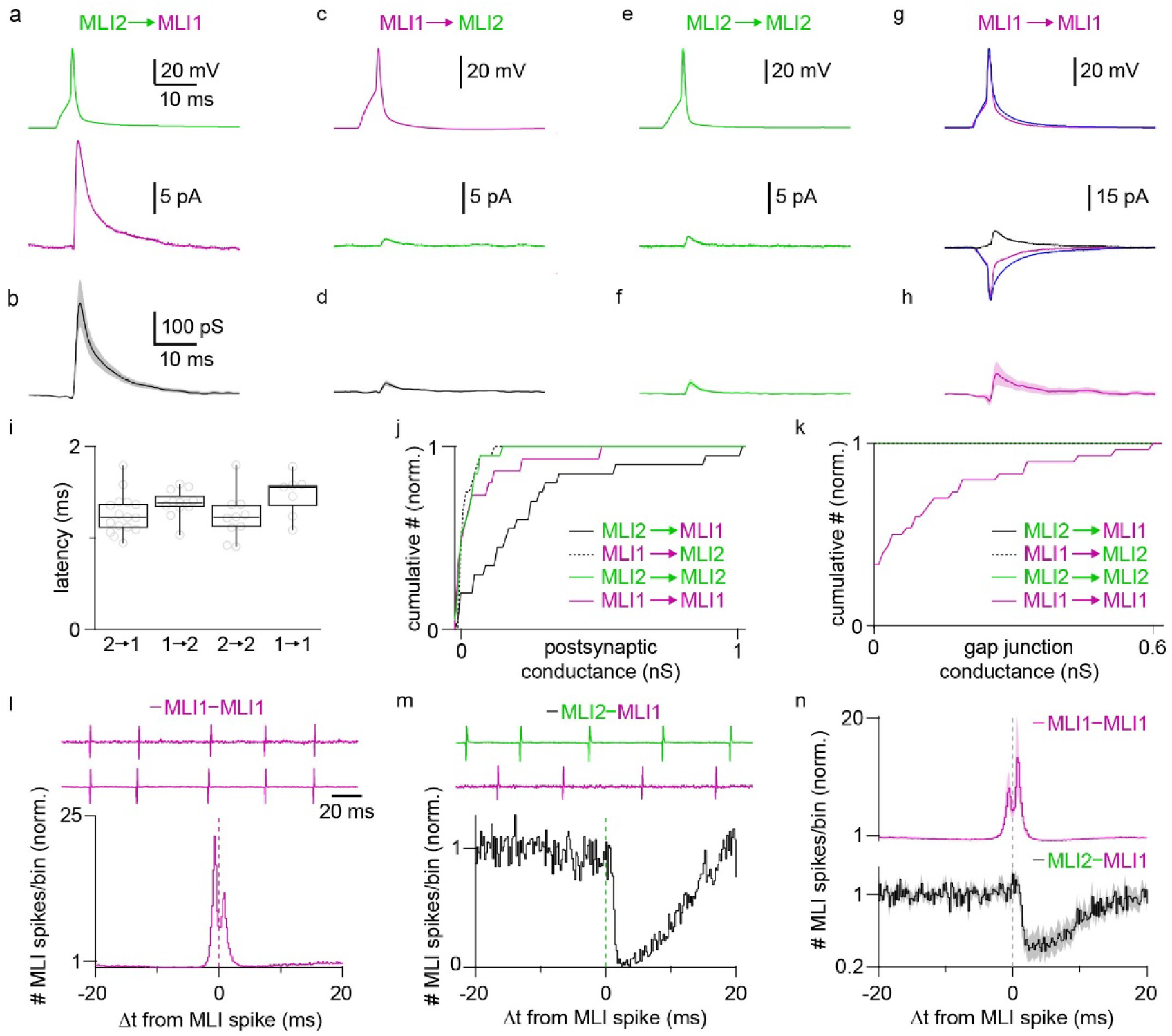
Synaptic connections and electrical connections for MLI-MLI pairs. **a.** A paired recording with an action potential evoked in a presynaptic MLI2 (current clamp) and the resulting IPSC recorded in an MLI1 (voltage clamp). **b.** Average synaptic current for MLI2-MLI1 pairs (n=20, p=2E-08). **c, d).** Same as **a, b**, but for MLI1-MLI2 pairs (n=20, p=0.00037). **e, f).** Same as **a, b**, but for MLI2-MLI2 pairs (n=20, p=0.0029) **g, h)** Same as **a, b**, but for MLI1-MLI1 pairs (n=18, p=0.87). The GABA_A_R antagonist gabazine was washed in (*blue*) and the difference between responses evoked in the presence and absence of gabazine are shown (*black*). **i.** Summary of latencies for MLI-MLI connections. **j.** Normalized cumulative plot of inhibitory conductances for all pairs of MLIs. **k.** Normalized cumulative plot of gap junction conductances for all pairs of MLIs. **l.** Cross-correlogram of an MLI1-MLI1 pair with on-cell recordings shown above. **m.** As in **l**, but for an MLI2-MLI1 pair. **n.** Average cross correlograms for MLI1-MLI1 and MLI2-MLI1 pairs. Synchrony was measured as the average normalized spike count from -1 ms to +1 ms (MLI1-MLI1 n=7, p=6E-04; MLI2-MLI1 n=8, p=0.56).

Inhibition was present in 39% (7/18) of the MLI1-MLI1 pairs (**Fig. 2j**), but the average conductance was small (50.3 ± 32.5 pS, **Fig. 2h**). The short latencies of all MLI-MLI synaptic connections suggests that they are all direct (**Fig. 2i**). A comparison of the cumulative plots as a function of connection strength (**Fig. 2j**) shows that synapses between all types of MLIs are present, but that MLI2-MLI1 synaptic connections are the most prevalent, and the largest. This is consistent with MLI2s disinhibiting PCs by suppressing MLI1 firing.

Electrical coupling has been shown to promote synchronous firing on the millisecond time scale in many types of neurons^30–32^, including MLIs^6,7,33^, but the observation that only MLI1-MLI1 pairs are electrically coupled (**Fig. 2k**) suggests that synchronous firing might be restricted to MLI1-MLI1 pairs. We tested this possibility by recording spontaneous spiking from identified MLI-MLI pairs, and found that 6/7 MLI1- MLI1 pairs and 0/8 MLI1-MLI2 pairs fired synchronously on the millisecond time scale (**Fig. 2ln**), whereas 7/8 MLI2s transiently suppressed the firing of nearby MLI1s (**Fig. 2mn**).

## Synaptic targets from EM reconstructions

We also used a large-scale EM dataset^34^ and serial EM reconstructions to assess MLI connectivity. The initial challenge was to identify MLI subtypes. Fortunately, we found that the cell bodies of MLI1 and MLI2 had distinctive morphological features. This is shown for fluorescent images of an electrophysiologically identified MLI1 and MLI2 (**Fig. 3a**, *left*), and for EM reconstructions of two MLIs (**Fig. 3a**, *right*). Spines are present on the cell bodies and proximal dendrites of MLI1s, whereas the cell bodies and proximal dendrites of MLI2s are smooth. We analyzed 179 MLIs (**Fig. 3b**): 139 MLIs with spiny cell bodies and proximal dendrites (**Fig. 3b**, *purple*), and 40 smooth MLIs (**Fig. 3b**, *green*) were distributed throughout the molecular layer. The ratio of spiny to smooth MLIs was 3.5:1 (139:40), which is comparable to the 3.1:1 ratio based on snRNA-seq data (32,716 MLI1s: 10,608 MLI2s)^2^. We fully reconstructed 30 spiny MLIs and 15 smooth MLIs. Smooth MLIs did not have collaterals that contributed to pinceaux (0/15), and all spiny MLI1s whose cell bodies were within 100 µm of the PC layer (18/18) had collaterals that contributed to pinceaux (**Fig. 3e, Extended Data Fig. 4**). These findings are similar to the morphologies of electrophysiologically identified MLI1s and MLI2s (**Fig. 1. Extended Data Fig. 2ij**).

**Figure 3.**
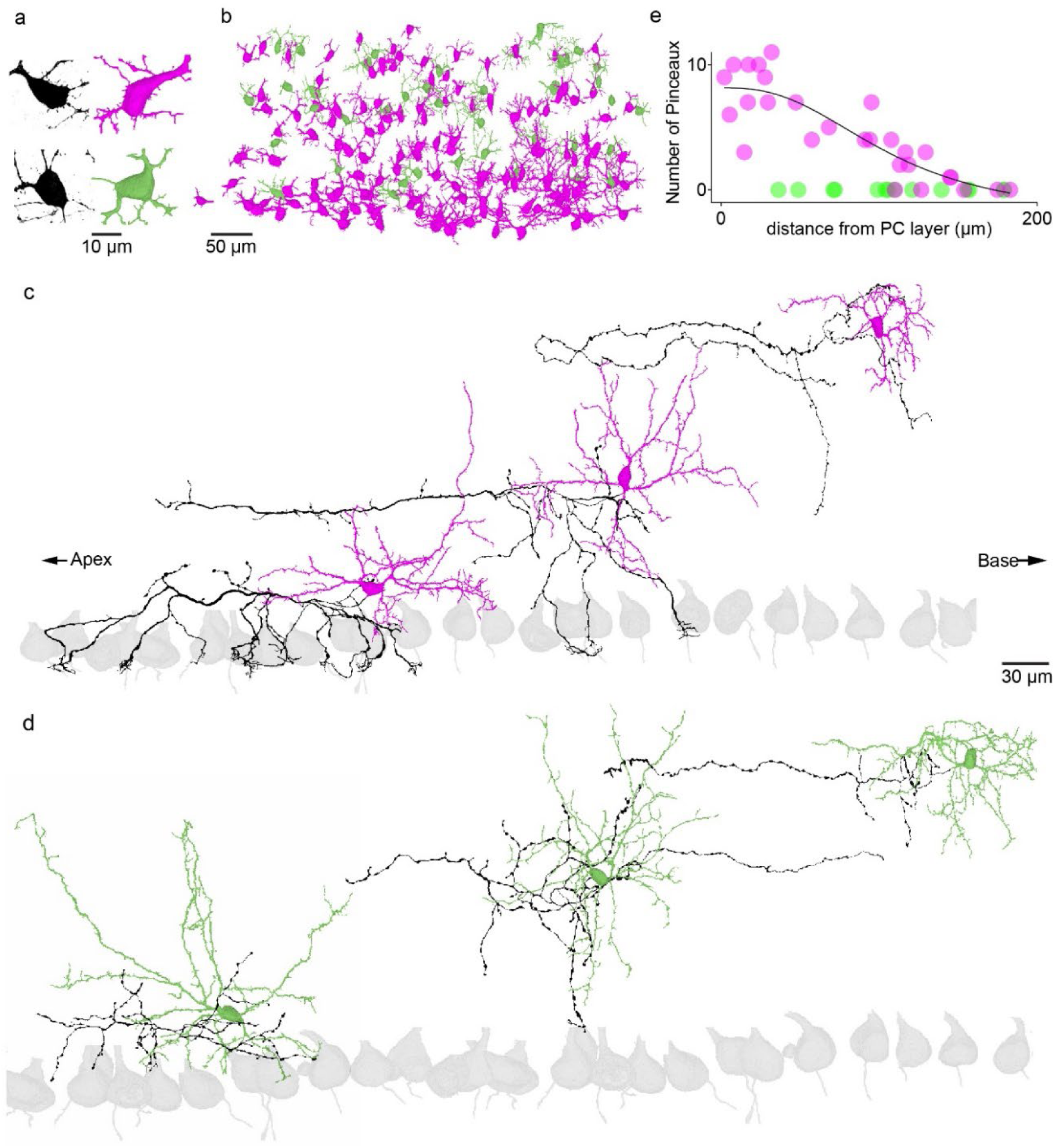
Serial EM reconstructions of MLIs. **a.** (*top left*) Image of the cell body of a fluorescently labelled MLI1. (*top right*) EM reconstruction of an MLI with prominent spines on the soma and proximal dendrites (MLI1, *purple*). (*bottom left*) Image of the cell body of a fluorescently labelled MLI2. (*top right*) EM reconstruction of a smooth MLI (MLI2, *green*). **b.** Cell bodies of EM reconstructed spiny (MLI1, *purple*)) and smooth (MLI2, *green*) MLIs. **c.** Reconstructed spiny MLIs (MLI1s) in different positions in the molecular layer, with dendrites (*purple*), and axons (*dark purple*) shown. **d.** Reconstructed smooth MLIs (MLI2s) in different positions in the molecular layer, with dendrites (*green*), and axons (*dark green*) shown. **e.** Summary of the number of pinceaux for MLIs at different distances from the PC layer.

Based on the appearance of cell bodies, the correspondence of the spiny/smooth ratio to the MLI1/MLI2 ratio observed previously, and the similarity of axonal morphologies for fluorescently labelled MLI1s (**Fig. 1ab**) and reconstructed spiny MLIs (**Fig. 3c**), we conclude that spiny MLIs are MLI1s and smooth MLIs are MLI2s.

Reconstructions of MLI1s are shown for neurons located in the lower, middle and upper molecular layer, with PC somata shown in grey (**Fig. 3c, Extended Data Fig. 4**). The lowest MLI1 has a typical basket cell morphology, with 14 axon collaterals extending to the initial segments of 11 PC axons. The middle MLI1 extended an axon for 300 µm within a sagittal plane to form a large number of synapses, and also extended collaterals that contributed to three pinceaux. The upper MLI1 had a classical stellate cell appearance and did not contribute to any pinceaux. MLI1s contributed to pinceaux in a graded manner that depended upon the position in the molecular layer (**Fig. 3e**), which is consistent with the observation that MLI1s have molecular properties that continuously vary with distance from the PC layer^2^. In contrast, all MLI2s, including cells located near the PC layer, had classical stellate cell morphologies, and did not extend axon collaterals below the PC layer to contribute to pinceaux (**Fig. 3de, Extended Data Fig. 4**). These results overturn the long-standing view that all MLIs near the PC layer are basket cells.

We reconstructed ten MLI1s (**Fig. 4a**) and ten MLI2s (**Fig. 4b**) and their targets (**Extended Data Fig. 5**, **Extended Data Fig. 6**), to quantify the output synapses of MLI1s and MLI2s. MLI1s primarily synapsed onto PCs (**Fig. 4cd, Extended Data Fig. 7a**), and MLI2s primarily synapsed onto MLI1s (**Fig. 4ef, Extended Data Fig. 7b**). The spatial locations of synapses made by MLI1s are summarized by displaying the synapse locations relative to the somata (**Fig. 4c**). Most synapses made by MLI1s were located within 200 μm of the somata. Synapses made by MLI2s onto MLI1s were more spatially restricted, and were present at higher densities towards the apex of the lobule and towards the PC layer (**Fig. 4e**). This arrangement of synapses is consistent with the tendency of MLIs to more strongly inhibit other MLIs below them within the molecular layer^5^, but the preferential inhibition of MLI1s towards the apex of the lobule was unexpected, and suggests an interesting spatial component of disinhibition that is not yet understood. The MLI1s made more total synapses, more synapses onto PCs, fewer synapses onto MLI2s and fewer synapses onto MLI1s (**Fig. 4g**, *upper*). Individual MLI1s synapsed onto approximately the same number of neurons as MLI2s, synapsed onto more PCs, fewer MLI1s and fewer MLI2s (**Fig. 4g**, *lower*).

**Figure 4.**
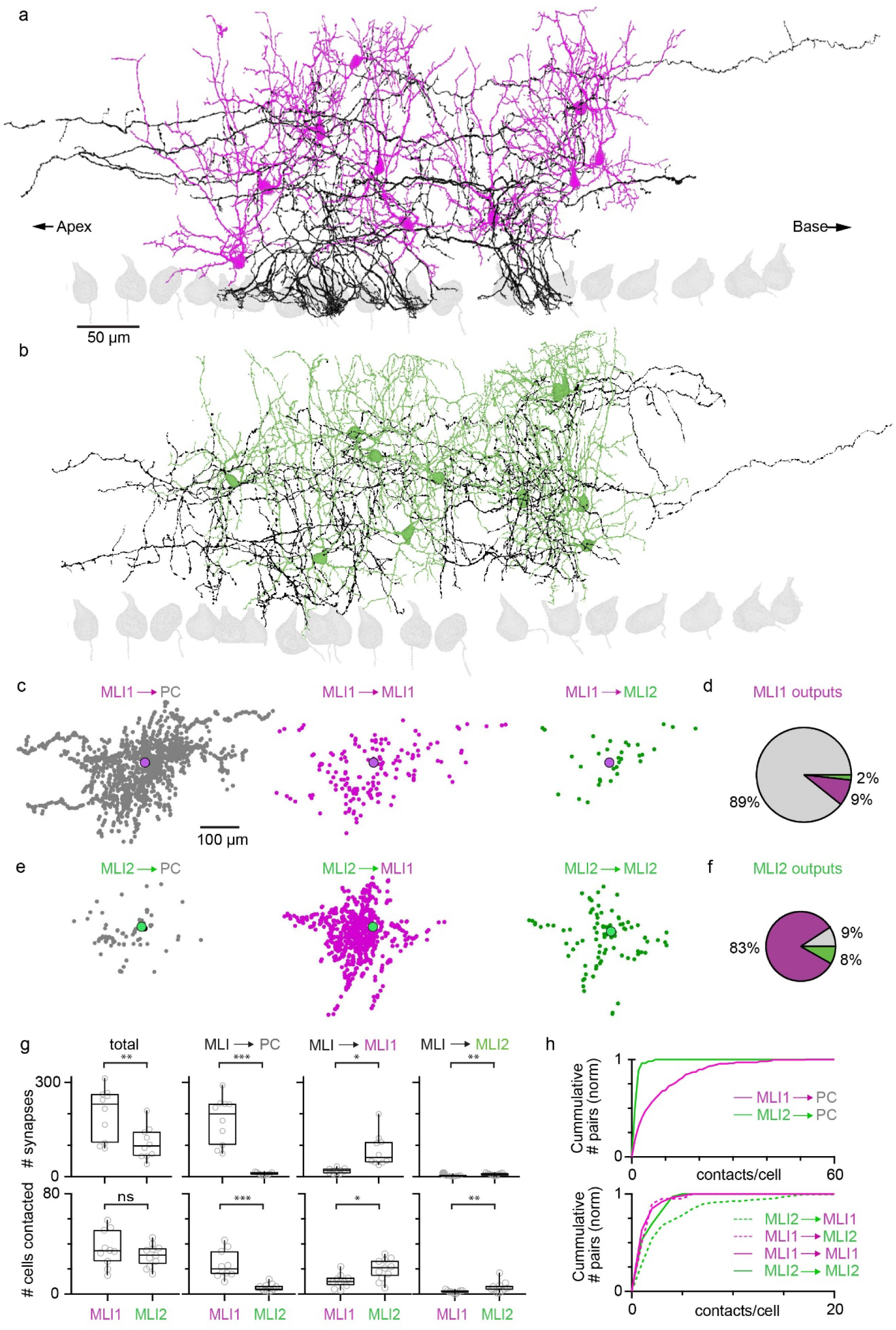
Target-dependence of synaptic connections made by MLI1s and MLI2s determined with serial EM reconstructions. **a.** Image of 10 reconstructed MLI1s. **b.** Image of 10 reconstructed MLI2s. **c.** (*left*) Positions of synaptic contacts made by MLI1s onto PCs relative to the MLI1 cell body. (*middle*) Positions of MLI1 to MLI1 synapses. (*right*) Positions of MLI1 to MLI2 synapses. **d.** Pie chart summarizing synaptic connections made by MLI1s onto different targets. **e.** As in (**c**) but for MLI2 synapses. **f.** As in (**d**) but for MLI2 synapses. **g.** (*top*) Summary of the total number of synapses and the number of synapses onto each type of target made by each MLI and MLI2. (*bottom*) Summary of the total number of contacted and the number of cells targeted by each MLI and MLI2. **h.** (*top*) Normalized cumulative plots of the number of synaptic contacts made by individual MLI1s and MLI2s onto PCs. (*bottom*) Normalized cumulative plots of the number of synaptic contacts made by individual MLI1s and MLI2s onto different types of MLIs.

We also quantified the number of contacts made by MLIs onto individual cells of each type (**Fig. 4h**). MLI1s made many synapses onto each PC, but there was a wide range in contacts per cell. MLI2-PC connections consisted of a small number of contacts, which is consistent with the connection strengths observed in **Fig. 1**. MLI2s primarily contacted MLI1s, and there was a wide range of connections per cell, whereas MLI→MLI2, MLI1→MLI1 and MLI2→MLI2 connections had very few contacts, in agreement with the connection strengths observed in paired recordings (**Fig. 2**). On average MLI1s made 202 synapses onto 38 cells, with 179 contacts onto an average of 24 PCs, and MLI2s made 107 synapses onto 31 cells, with 84 contacts onto 20 MLI1s. These reconstructions suggest that there is considerable variability in the number of contacts per cell, which is also consistent with our electrophysiological experiments. These results establish that MLI1 and MLI2 are distinct circuit elements with specialized connectivity, and the traditional view that all MLIs primarily inhibit PCs must be revised.

## MLI subtypes *in vivo*

Determining the activity of MLI1s and MLI2s during behavior promises to provide insight into their different functional roles. Multielectrode probes make it possible to simultaneously record from many MLIs and PCs, but discriminating between MLI1s and MLI2s *in vivo* is challenging. Although the *Nxph1^Cre^* line that we made is useful for slice recordings, it is not sufficiently selective to optically tag MLI2s for *in vivo* identification or for optogenetic studies. Nonetheless, our slice and EM studies make testable predictions regarding the properties of MLI1s and MLI2s *in vivo* that can help identify MLI1s and MLI2s in single unit recordings (**Fig. 5a**).

**Figure 5.**
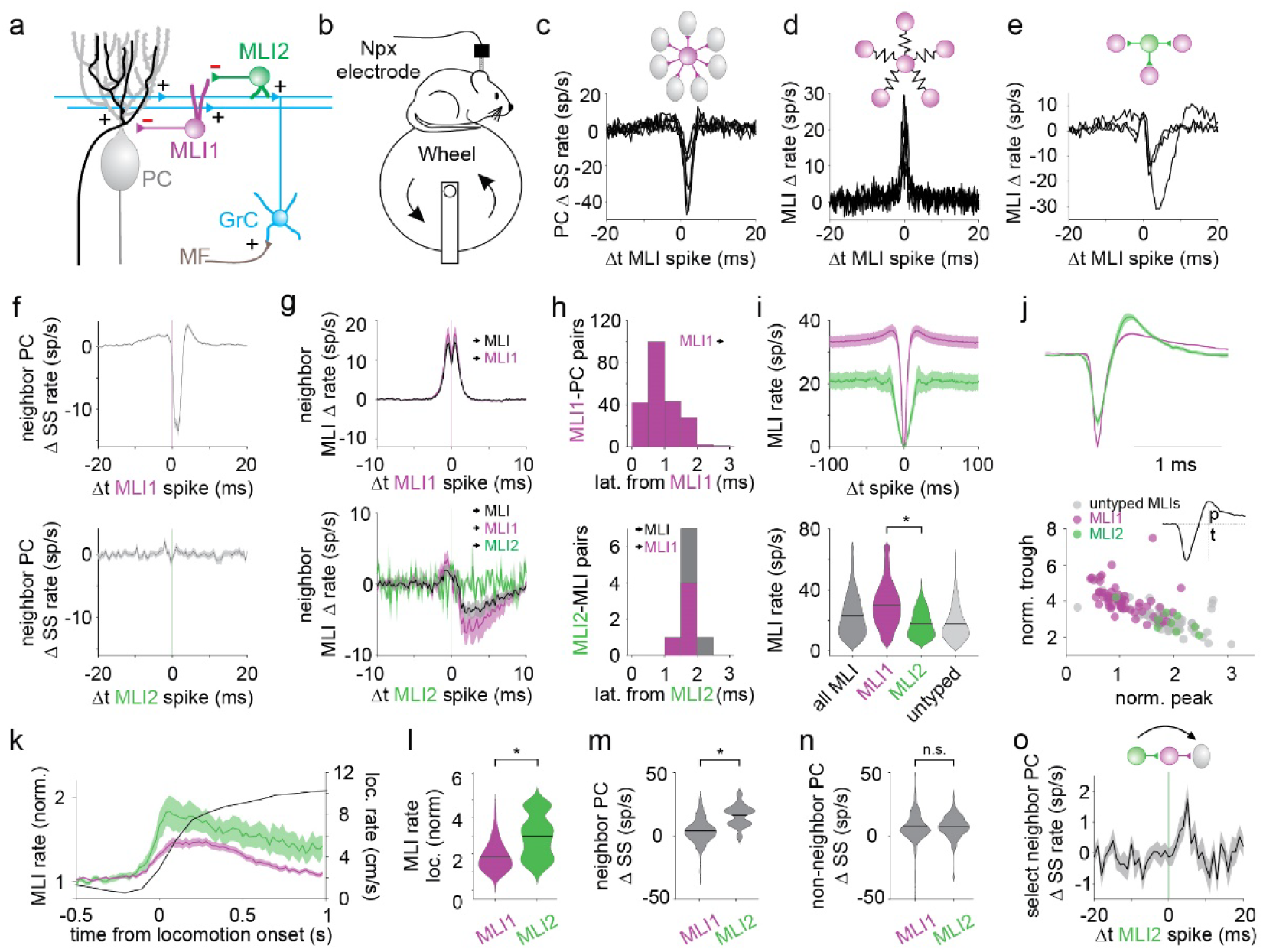
*In vivo* recordings of MLI activity in behaving mice. **a.** Schematic of the cerebellar cortex, including MLI1 and MLI2. **b.** Neuropixels probes were used to record single-unit activity from the cerebellar cortex of awake mice head-fixed on a freely moving wheel and the properties of neighboring MLIs and PCs (<125 μm separation) were analyzed. **c.** An MLI inhibits seven PCs. **d.** An MLI fires synchronously with six MLIs. **e.** An MLI inhibits 3 MLIs. **f.** PCs are inhibited by putative MLI1s but not by putative MLI2s (338 putMLI1-PC pairs; 19 putMLI2-PC pairs). **g.** Putative MLI1s fire synchronously with other nearby MLIs (top, 335 MLI1-MLI pairs and 286 MLI1- MLI1 pairs). Putative MLI2s inhibit other MLIs (bottom, 32 MLI2-MLI pairs, 15 MLI2-MLI1 pairs, and 2 MLI2-MLI2 pairs). **h.** Latencies of putative MLI1-PC inhibition (top) and from MLI2s onto MLIs (bottom). MLI1, pink; all MLIs, gray. **i.** Autocorrelograms and firing rates during quiescence for all MLIs, MLI1s, MLI2s, and unclassified MLIs (p = 0.012, 59 putative MLI1s and 9 putative MLI2s). **j.** Putative MLI1 and MLI2 mean waveforms (top) and distributions of waveform trough and peak sizes for individual MLIs. (bottom). **k.** Mean PSTHs of MLI1s and MLI2s aligned to locomotion onset (*black trace*). **l.** Mean firing rate changes of MLI1s and MLI2s during the first 100 ms of locomotion normalized to firing rates during quiescence (p = 0.027). **m.** PC firing rates during periods segmented according to the highest and lowest firing rates of neighboring MLIs of each type (p = 0.00002, MLI1: n = 268 pairs, MLI2: n = 17 pairs). **n.** PC firing rates during periods segmented according to the highest and lowest firing rates of non- neighboring MLIs of each type (p = 0.36, MLI1: n = 674 pairs, MLI2: n = 89 pairs). **o.** Increase in PC spike rate after MLI2 spikes for simultaneously recorded MLI2-MLI1-PC connections (16 pairs).

We recorded using Neuropixels probes from awake mice head-fixed on a freely moving wheel (**Fig. 5b, Extended Data Fig. 8**), and identified 110 PCs based on characteristic complex spike responses (**Extended Data Fig. 8b**) and 132 MLIs by their firing rates (>3Hz) and their laminar location (**Extended Data Fig. 8cd**). Based on an MLI1:MLI2 ratio of 3.5:1, it is estimated that more than 100 of these are MLI1s and the rest are MLI2s. For cross correlograms between nearby MLIs and PCs, 79 MLIs produced a strong (Z-score>4), short latency (<3 ms) decrease in firing rates in at least one nearby PC (**Fig. 5c**). We classified these MLIs as putative MLI1s based on our finding that MLI1-PC synapses are much stronger than MLI2-PC synapses. On average, putative MLI1s were in close proximity (<125 µm) to 4.3 PCs, and inhibited 3.4 of them. Putative MLI1 inhibition of PCs decreased target PC firing by an average of 19.1 ± 0.7 sp/s for 1.30 ± 0.06 ms (**Extended Data Fig. 8e**) with a latency of 0.70 ± 0.04 ms (**Fig. 5h**). Of the 72 putative MLI1s that were close (<125 μm) to at least one other MLI, 59 fired synchronously on the millisecond time scale with at least one other MLI (**Fig. 5d**), and these MLI-MLI cross correlograms (**Fig. 5g**) are similar to those observed for MLI1-MLI1 pairs in brain slices (**Fig. 2ln**). On average, putative MLI1s fired synchronously with 1.7 of 4.3 neighboring MLIs (mean increase of 19.0 ± 1.6 sp/s for 3.4 ± 0.3 ms, **Extended Data Fig. 8**). Of 335 putative MLI1-MLI pairs, 130 fired synchronously on the millisecond time scale, and only five were inhibitory. These findings indicate that a large fraction of MLI1s fire synchronously with each other in awake mice, and that they have the capacity to transiently pause the firing of multiple PCs, which could evoke precisely timed spikes in cerebellar nuclei projection neurons^18^.

As expected, MLI2s were less numerous and more difficult to identify *in vivo*. We found four MLIs that had at least three nearby PCs and did not inhibit any of them, and six additional MLIs that inhibited other MLIs, but no PCs. One of these fired synchronously with another MLI. This left nine putative MLI2s, seven of which inhibited other MLIs, as in **Fig. 5e**. Average cross correlograms indicate that putative MLI2s did not inhibit PCs (**Fig. 5f**), as required by our selection criteria, but they inhibited nearby MLIs (**Fig. 5g**). Putative MLI2s inhibited 12/32 nearby MLIs and decreased MLI firing by 11.7 ± 2.2 sp/s for 4.0 ± 0.9 ms (**Fig. 5jk**) with a short latency (**Fig. 5h**). Average baseline firing rates were higher for MLI1s than MLI2s, but there was some overlap in the firing rates for the two types of MLIs (**Fig. 5i**).

These observations are consistent with the MLI1 and MLI2 spontaneous firing rates in brain slice^2^. We also found that the average MLI1 and MLI2 waveforms differed, although overlap in the waveform properties indicated that for individual cells the waveform alone cannot be used to discriminate between MLI1 and MLI2 (**Fig. 5j**).

At the onset of locomotion, the firing rates of both MLI1s and MLI2s increased (**Fig. 5k,l**). However, MLI2s showed larger increases in firing (**Fig. 5l**), consistent with their enhanced excitability as described *in vitro*^2^. These recordings also allow us to assess the relationships between putative MLI1 and MLI2 firing rates, and PC firing rates. We determined the 100 ms intervals when MLI1s fired fastest and slowest, and measured the difference in nearby PC firing rates for these conditions (PC_MLI1fast_ – PC_MLI1slow_). We repeated this calculation for MLI2 firing (PC_MLI2fast_ – PC_MLI2slow_). This analysis showed that for nearby PCs, PC_MLI1fast_ – PC_MLI1slow_ was slightly elevated (**Fig. 5m**), reflecting the general trend of locomotion to elevate PC, MLI1 and MLI2 firing (**Fig. 5k**). However, PC_MLI2fast_ – PC_MLI2slow_ was elevated to a much larger extent than PC_MLI1fast_ – PC_MLI1slow_, consistent with ML2s disinhibiting PCs *in vivo* (PC = 3.88 ± 0.60 sp/s grouped by MLI1 activity, ΔPC = 16.1 ± 2.0 sp/s grouped by MLI2 activity, **Fig. 5m**). As a control, we repeated this for MLI-PC pairs with >125 μm separation, and there was no difference between PC_MLI1fast_ – PC_MLI1slow_ and PC_MLI2fast_ – PC_MLI2slow_ (**Fig. 5n**). Although the average MLI2-PC cross correlogram did not reveal disinhibition following single spikes (**Fig. 5f**), we were able to examine rare cases where we simultaneously recorded from an MLI2 that inhibited a nearby MLI1 that in turn inhibited a PC. The average MLI2-PC cross correlogram for such conditions showed that a single spike in a putative MLI2 transiently elevated firing in nearby PCs with a time course consistent with disinhibition mediated by a disynaptic connection (**Fig. 5o**). These findings suggest that our criteria for selecting MLI subtypes allows us to identify MLI1s and MLI2s, and that MLI2s have a characteristic motif that allows them to transiently disinhibit PCs *in vivo*, whereas MLI1s fire synchronously to transiently suppress PC firing.

## Discussion

We used paired recordings, serial EM reconstructions and *in vivo* recordings in behaving animals to establish that MLI subtypes are specialized to serve distinct roles. MLI1s primarily inhibit PCs, they are electrically-coupled to each other, they can fire synchronously on the millisecond time scale, and they can synchronously inhibit PCs. MLI2s are not electrically coupled to each other, they primarily inhibit MLI1s, and they disinhibit PCs. This establishes that together MLI subtypes comprise a disinhibitory circuit that expands the computational potential of the cerebellar cortex. The specialized circuit properties of MLI subtypes described here indicate that our view of cerebellar processing and models of the cerebellar cortex require a major revision. Models of the cerebellar cortex have generally only considered MLI inhibition of PCs and have not considered MLI-MLI inhibition or electrical coupling of MLIs^35–37^. Our findings establish that, to understand cerebellar processing, and how cerebellar plasticity is regulated, it is necessary to consider MLI1 synchrony, synchronous pauses in PC firing, and MLI2 disinhibition of PCs.

The discovery that MLI2s are disinhibitory interneurons indicates that the molecular layer of the cerebellum shares a circuit motif that plays an important processing role in many brain regions, including layer 1 of the cerebral cortex^9–14^. Whereas elevated MLI activity suppresses calcium signaling in PC dendrites and suppresses the induction of LTD at granule cell to PC synapses^19,20^, the finding that MLI2s inhibit MLI1s suggests that it is possible to bidirectionally influence LTD induction. It seems likely that MLI2s provide the primary source of disinhibition that gates learning in the cerebellar cortex^8^. More generally, MLI2s provide a means of countering granule cell excitation of MLI1s to keep them in a responsive range, and they allow bidirectional regulation of MLI1 firing rates. This also allows flexible and bidirectional regulation of PC firing rates in a manner that could not be readily achieved with a simple feedforward inhibitory circuit where inhibition scales with incoming excitation. The relative simplicity of the cerebellar cortex, and our ability to characterize the granule cell, MLI1, MLI2 and PC synapses, promises to lead to new insights into the advantages of disinhibition.

Our paired recordings in slices established that MLI1-MLI1 pairs are electrically coupled and fire synchronously with each other on the millisecond time scale, but that other combinations of MLIs are not electrically coupled with each other and do not fire synchronously. The putative MLI1-MLI1 cross-correlograms we observed *in vivo* are very similar to those seen for electrically-coupled MLI1-MLI1 pairs in brain slices, suggesting that electrical coupling in MLI1s underlies their synchronous firing *in vivo*.

Electrical coupling also allows MLI1s to share charge from synaptic inputs, as has been described for Golgi cells^38^, and consequently MLI1s are not completely independent circuit elements. EM reconstructions indicate that direct electrical coupling is only possible for MLI1s whose dendrites reside within approximately the same parasagittal plane, making electrical coupling between MLI1s suited to coordinating firing within parasagittal microzones^39–41^. Transient decreases in PC firing arising from synchronous inhibition from multiple MLI1s are suited to promote precisely-timed increases in firing within cerebellar nuclei^18^. This suggests that MLI1-induced synchronous pauses in PC firing could help to gate the output of the cerebellar cortex.

## Methods

### Animals

Animal procedures were performed in accordance with the NIH and Animal Care and Use Committee (IACUC) guidelines and protocols approved by the Harvard Medical School Standing Committee on Animals. C57BL/6 mice were obtained from Charles River Laboratories. *Gjd2-EGFP* mice were obtained from MMRRC (stock # 030611-UCD)^29^. *Nxph1^Cre^* mice were crossed with *Ai14* reporter mice (Jackson Labs, stock # 007908)^42^ and kept in a mixed genetic background. Animals of either sex were randomly selected for experiments. Animals were housed on a normal light–dark cycle with an ambient temperature of 18–23 °C with 40–60% humidity.

Mice for *in vivo* recordings (15 mice, 8 male, >P55) with a C57 or CBA background (1 C57, 2 C57 x ckit, 12 C57 x CBA) were used in accordance with approval from the Duke University Animal Care and Use Committee. Animals were housed on a normal light–dark cycle, and animals of either sex were randomly selected for experiments.

#### Generation of Nxph1^Cre^ mice

Easi-CRISPR ^43^ was used to generate the *Nxph1^Cre^* mouse line, resulting in the insertion of a p2a-Cre recombinase after the *Nxph1* exon 3 stop codon, TGA (**Extended Data Fig. 1a**). First, CAS9 (PNABio, CP01-50) sgRNA (CCUGUUCAUCUUCAUCCGGA, Synthego) and ssDNA (p2a-Cre cassette flanked by 150 nucleotides homology arms on 5’ and 3’ ends, IDT) were injected (Beth Israel Deaconess Medical Center Transgenic Core Facility) in fertilized eggs of FVB mice, then founders carrying the desired insertion were detected through PCR and subsequently sequenced (Biopolymers Facility at Harvard Medical School) around the insertion to confirm intact cassette sequence.

#### HCR-FISH and immunohistochemistry

Based on the selective expression of *Gjd2* in MLI1s and *Nxph1* in MLI2s^2^, we hoped that *Gjd2-EGFP* mice and *Nxph1^Cre^Ai14* mice would allow us to target MLI1s and MLI2s, respectively. We used HCR-FISH in combination with immunohistochemistry to assess the suitability of these mouse lines. Acute cerebellar slices (1 midline slice per mouse) from p28-p45 mice were prepared as described, and fixed for 2 hours in 4% paraformaldehyde in PBS (Biotum) at 4 °C. Slices were stored overnight in 70% ethanol in RNase-free water at 4 °C. A floating slice HCR protocol^2^ was performed with the following probes and matching hairpins (Molecular Instruments): sortilin related VPS10 domain containing receptor 3 (*Sorcs3*), and neurexophilin 1 (*Nxph1*). MLI1s express *Sorcs3* and *Gjd2*, MLI2s express *Nxph1*. Amplification hairpins were B1-647 (Alexa 647) and B2-488 (Alexa 488) or B2-594 (Alexa 594) for fluorescence imaging in conjunction with TdT or GFP. Anti-GFP immunohistochemistry was performed between permeabilization and hybridization. Slices were incubated in blocking solution containing primary antibody (chicken anti-GFP, Abcam, 1:1000) overnight at room temperature. Slices were washed in 2x SSC (3 x 5 min) and incubated in blocking solution containing secondary antibody (anti-chicken Alexa 488, Abcam, 1:1500) for 2 hours at room temperature. Slices were washed in 2x SSC (3 x 5 min), postfixed in 4% paraformaldehyde for 10 min, and washed in 2x SSC (3 x 5 min) before hybridization.

Slices were mounted on slides (Superfrost Plus, VWR) with mounting medium (Fluoromount, ThermoFisher) and no.1 coverslips. Images were acquired with a Leica Stellaris X5 confocal microscope using a 63x oil immersion objective (1.4 NA, Olympus). The reporter and HCR probe/hairpin channels were imaged with 180 nm resolution in a 10-20-µm thick, 0.5-µm interval tiled z series in lobule IV/V. Noise was reduced using a median filter with a 2-pixel radius for each focal plane in Fiji (ImageJ). *Sorcs3*+ and *Nxph1*+ cell locations in the molecular layer were manually labelled using the multi-point tool in Fiji. TdT or GFP fluorescence in each cell was averaged within a 7-µm diameter circular mask in Matlab (Mathworks).

TdT labelling was observed in MLIs in *Nxph1^Cre^Ai14* mice (**Extended Data Fig. 1b**). Essentially all MLI2s (*Nxph1*+/*Sorcs3*- cells in the molecular layer) were labelled, and some were very intensely labelled (**Extended Data Fig. 1b-e**). Approximately 55% of the MLI1s (*Nxph1*-/*Sorcs3*+ cells in the molecular layer) had very low fluorescence levels, and the rest had moderate fluorescence levels. We found that targeting bright cells in *Nxph1^Cre^Ai14* mice allowed us to target MLI2s (although we also performed a complete electrophysiological characterization to insure the identity of MLI subtypes). Although *Nxph1^Cre^*is useful for identifying MLI2s, it does not provide sufficient selectivity for optogenetic activation or suppression of MLI2s.

We characterized *Gjd2-EGFP* mice in a similar manner, and found many MLIs were labelled in these mice (**Extended Data Fig. 1fg**). Quantification of the GFP fluorescence intensity showed the most MLI2s had low fluorescence levels, but that some were moderately fluorescent. Conversely, some MLI1s had moderate fluorescence levels, but some were very bright. Almost all bright cells were MLI1s. Ultimately, it was not necessary to use *Gjd2-EGFP* mice to identify MLI1s, because most MLIs are MLI1s, and electrophysiological characterization of cells allowed us to readily establish that a cell was an MLI1.

### Slice electrophysiology

#### Slice preparation

Acute parasagittal slices (230-μm thick) were prepared from p28-45 C57BL/6, *Gjd2-EGFP*, or *Nxph1^Cre^Ai14* mice. Mice were anaesthetized with an intraperitoneal injection of ketamine (10 mg kg^−1^) and perfused transcardially with an ice-cold solution containing (in mM): 110 choline chloride, 7 MgCl_2_, 2.5 KCl, 1.25 NaH_2_PO_4_, 0.5 CaCl_2_, 25 glucose, 11.5 sodium ascorbate, 3 sodium pyruvate, 25 NaHCO_3_, equilibrated with 95% O_2_ and 5% CO_2_. Slices were cut in the same solution, and then transferred to artificial cerebrospinal fluid (ACSF) containing (in mM) 125 NaCl, 26 NaHCO_3_, 1.25 NaH_2_PO_4_, 2.5 KCl, 1 MgCl_2_, 1.5 CaCl_2_ and 25 glucose equilibrated with 95% O_2_ and 5% CO_2_ at 34 °C for 30 min. Slices were kept at room temperature until recording.

#### Recordings

MLI-PC and MLI-MLI paired recordings were performed at 32 °C with an internal solution containing (in mM): 150 K-gluconate, 3 KCl, 10 HEPES, 3 MgATP, 0.5 GTP, 0.5 EGTA, 5 phosphocreatine-tris_2_ and 5 phosphocreatine-Na_2_ (pH adjusted to 7.2 with KOH, osmolarity adjusted to 310 mOsm kg^−1^). Biocytin (0.2-1%) and Alexa 488 (0.1 mM) were added to the internal solution for MLIs and PCs, respectively. A calculated junction potential of -16.9 mV was corrected. Visually guided whole-cell recordings were obtained with patch pipettes of ∼1-3-MΩ resistance for PCs and ∼3-6-MΩ resistance for MLIs pulled from borosilicate capillary glass (BF150-86-10, Sutter Instrument). Slice recordings with PC leak currents greater than 500 pA were rejected. Electrophysiology data were acquired using a Multiclamp 700A or 700B amplifier (Axon Instruments), digitized at 20 kHz and filtered at 4 kHz. A subset of recordings was digitized at 100 kHz and down sampled to 20 kHz. Acquisition and analysis of slice electrophysiological data were performed using custom routines written in Igor Pro (Wavemetrics) and Matlab. The following receptor antagonists were added to the ACSF solution to block glutamatergic and glycinergic synaptic currents (in μM): 2 (R)-CPP, 5 NBQX, 1 strychnine. The GABA_A_R antagonist SR95531 (gabazine, 10 μM) was washed-in for a subset of experiments. All drugs were purchased from Abcam and Tocris.

We recoded from MLIs in the inner two-thirds of the molecular layer and determined the identity of MLI1s and MLI2s by characterizing a number of characteristic electrical properties (**Extended Data Fig. 2**). It was previously shown that spikelets are present in MLI1s but not in MLI2s, the input resistance is lower in MLI1s, and I_h_ is larger in MLI2s^2^. MLIs were held at −65 mV in voltage clamp for 30 s to determine if spikelets were present. Recordings were performed deep in slices because the spikelet detection relies on the activity of connected cells. Input resistances (*R*_i_) were determined using a 10 pA, 50 ms hyperpolarizing current step averaged over 50 trials. To activate the hyperpolarization-evoked currents (I_h_), MLIs were held at −65 mV and a 30 pA hyperpolarizing current step of 500 ms duration was injected. The amplitude of I_h_ was calculated as the difference between the maximal current evoked by the hyperpolarizing current step and the average steady-state current at the end (480–500 ms) of the current step. We also found that following action potential stimulation, the membrane potential decayed with a different time course in MLI1s and MLI2s. We imaged MLI morphology in a subset of MLIs identified on the basis of their electrical properties (see below), and found that 10/10 MLI1s and 0/6 MLI2s had collaterals that extended below the PC cell body layer to contribute to pinceaux. Some experiments were performed using *Nxph1^Cre^Ai14* mice to select MLI2s based on bright TdT fluorescence (**Extended Data Fig. 1bc**). We then confirmed MLI2 identity with electrical characterization.

To characterize synapses made by MLI1 and MLI2, presynaptic MLI spikes were evoked in whole-cell current clamp with 5 ms current injections at an average of 5 stimuli/s. ISIs were varied using a Gaussian distribution with standard deviation of 25% to prevent entraining firing in spontaneously firing populations of electrically-coupled MLIs. Presynaptic spontaneous MLI firing was suppressed by negative current injection. Postsynaptic responses were recorded in whole-cell mode in voltage clamp at -65 mV for 500 trials. Responses were also recorded at a holding potential of -30 mV for a subset of experiments. All synaptic currents are averages of 500 trials. Spontaneous action potentials from MLI-MLI pairs (within 5 µm of each other in the sagittal plane) were recorded in loose-patch configuration with ACSF-filled electrodes or in current clamp for 5-10 min.

#### Analysis

Postsynaptic currents were time-locked to the peak of the first derivative of presynaptic evoked spikes and low-pass filtered at 500 Hz. The amplitudes of outward and inward currents were measured as the average of 2-6 ms and 10-15 ms following the evoked spike, respectively (**Extended Data Fig. 3ab**). Spontaneous and evoked inhibitory events were detected on the first derivative of PC recordings filtered at 200 Hz, with a threshold of 1.5x the standard deviation. The events were integrated, and spontaneous events were subtracted using a linear fit over the 200 ms window before evoked spike onset (**Extended Data Fig. 3hi)**. The remaining change in events was measured as the average of 10-15 ms following the evoked spike (**Extended Data Fig. 3jk**). Responses were measured relative to baseline averaged 50 ms prior to the evoked spike, and amplitudes at baseline were measured 25 ms prior to the evoked spike. Pairs were determined to be connected if the response z-score was >2. Latency was measured as the half-max time for connected pairs. To calculate GABA_A_R-mediated synaptic currents in electrically coupled pairs, the average evoked postsynaptic response after gabazine wash-in was subtracted from the average evoked postsynaptic response before the wash-in. Gap junction conductance was calculated as g_j12_= 1/R_j_=ΔI_post_/ ΔV_pre_, with ΔI_post_ being the post-synaptic spikelet amplitude and ΔI_pre_ the pre-synaptic evoked spike amplitude^44^. To determine if MLIs fired synchronously, spontaneous action potentials were detected and manually verified for each cell, and cross correlograms and averages of the normalized spike count from -1 ms to 1 ms were calculated^33^.

#### Cell fills

Recorded MLIs and PCs were filled with 0.2-1% biocytin and 0.1 mM Alexa 488, respectively. Patch electrodes were retracted slowly until the cells resealed. Slices were transferred to a well-plate and submerged in 4% paraformaldehyde in PBS (Biotum). Spines were examined in dedicated experiments where MLIs were resealed immediately after cell type identification, within 5 min of recording. Slices were fixed overnight at 4 °C and kept in the dark to preserve Alexa fluorescence in PCs. Slices were washed in PBS (2 x 5 min) and incubated in 0.1% Tween in PBS containing streptavidin Alexa 594 conjugate (ThermoFisher, 1:1500) for 2 hours at room temperature. Slices were washed in PBS (3 x 5 min) and mounted on slides (Superfrost Plus, VWR) with mounting medium (Fluoromount, ThermoFisher) and no.1 coverslips.

Images of MLI-PC pairs were acquired with a Leica Stellaris X5 confocal microscope using a 63x oil immersion objective (1.4 NA, Olympus). The MLI (streptavidin-Alexa 594) and PC (Alexa 488) channels were imaged with 180 nm resolution in a tiled z series with a 0.5-µm interval. Noise was reduced in the PC channel using a median filter with a 4-pixel radius for each focal plane in Fiji (ImageJ). A maximum intensity z projection image was manually thresholded to create a binary image for each channel in Fiji. For MLI cell bodies examined for spines, images were acquired with a Leica SP8X confocal microscope using a 100x oil immersion objective (1.4 NA, Olympus). Cell bodies were imaged with 20 nm resolution and line averaging of 5 in a z series with a 100 nm interval, and the images were deconvolved using Hyugens software.

### Serial EM

We previously imaged and aligned a 770 μm X 750 μm X 53 μm volume of lobule V of the mouse cerebellum for EM reconstructions comprised of 1176 45-nm thick parasagittal sections^34^. We used automated image segmentation to generate neuron boundaries^34^. To identify MLI1 and MLI2 subtypes, cell bodies located in the molecular layer were reconstructed and the presence of spines on the soma and proximal dendrites was evaluated.

Interneurons with spiny somata were characterized as MLI1 and smooth somata were characterized as MLI2. Synaptic outputs of both cell types were analyzed using an artificial neural network that we previously trained to automatically detect synapses^34^. The network is implemented using a python package (NetworkX). Synapses were manually proofread to validate each postsynaptic target using MD-Seg^34^.

Synapses were identified using automated synapse detection and proofread manually for ten MLI1 and ten MLI2 cells. The cell bodies of these MLIs were located near the middle of the volume, and it was possible to reconstruct dendritic regions of the target cells and their somata to allow subtype determination of most target MLIs (2.3% of synapses were made onto MLIs that could not be categorized because the cell body of the target cell was not contained within the EM series). Synapses were identified by characteristic ultrastructural features of GABAergic synapses^45^, including a synaptic cleft with a flattening of apposed pre- and post-synaptic membranes and clustering of synaptic vesicles near the presynaptic specialization. Plots were rotated 5.62 degrees to compensate for tilt of the PC layer in the volume. Of the 2025 synapses made by the ten MLI1s, 19 were onto MLIs whose subtype could not be determined, 1 was onto a granule cell layer interneuron. Of the 1070 synapses made by the ten MLI2s, 50 were onto MLIs whose subtype could not be determined, 1 was onto a granule cell layer interneuron, and 1 was onto a candelabrum cell. We did not find any MLI1 or MLI2 to Golgi cell synapses, which is consistent with a previous electrophysiological/optogenetic study^46^.

To examine morphologies, 30 MLI1s and 15 MLI2s distributed throughout the molecular layer were reconstructed (**Fig 3e**). Axonal projections made below the Purkinje cell layer and proximal to the axon initial segment of a PC were identified as contributing to a pinceau. The number of pinceaux were plotted against the position of the MLI somata, and the relationship was visualized by applying a Hill equation fit to the data.

### *In vivo* recordings

#### Surgical Procedures

Animals underwent a headposting surgery weeks before recording, during which a titanium headpost (HE Palmer) was affixed to the skull and a stainless steel ground screw (F.S. Tools) was inserted over the left cerebellum, both secured with metabond (Parkell). Mice received dexamethasone (3 mg/k, subq) 4- 24 hours before surgery and an initial dose of ketamine/xylazine (50 mg/kg and 5 mg/kg, IP) and carprofen (5 mg/kg) 20 min before induction with isoflurane anesthesia. Isoflurane was administered at 1.0-2.0% throughout surgery to maintain appropriate breathing rates and prevent toe pinch response, which were monitored throughout the duration of the surgery. Body temperature was maintained with a heating pad (TC-111 CWE). Mice received bupranex and cefazolin (0.05 mg/kg and 50 mg/kg respectively, subq) twice daily for 48 h after surgery and were monitored daily for 4 days. After 2+ weeks of recovery, mice received dexamethasone (3 mg/k) 4-24 hours before recordings. Craniotomies (approx. 0.5-1.5 mm) were opened on the first day of recording over lobule simplex or Crus I, under 1- 2% isoflurane anesthesia, and were sealed between recordings using Qwik-Cast (WPI) covered by Metabond. Craniotomies could be re-opened for subsequent recordings under brief (<30 min) 1-2% isoflurane anesthesia.

#### In vivo electrophysiology

After recovery from headpost placement, mice were habituated to be head fixed on a freely moving wheel for at least 30 min over 3 days. Mice were given dexamethasone (2 mg/k) 4-24 hours before recording. After the craniotomy was opened, mice were head fixed on the wheel and allowed to recover from anesthesia. Neuropixels 1.0 electrodes were positioned in the right lateral cerebellum between 0-2 mm lateral and 6-7 mm posterior to Bregma. The electrode was lowered at rate of 1-5 μm/sec through the cerebellar cortex to reach a final placement of 1,000 – 2,500 microns into the cortex. Tissue was allowed to relax for 30 minutes and recordings lasted 30-90 min. Mouse movements were recorded with a rotatory encoder (YUMO) attached to the wheel and licking was monitored with an optical lick sensor (custom). In some experiments, mice were water deprived for 3-7 days before recording and received a water reward every 20-40 sec during recording to facilitate locomotion. All metrics were computed during quiescent periods unless otherwise noted. After the last day of recording, animals were deeply anesthetized with a ketamine/xylazine (350 mg/kg and 35 mg/kg, IP) and perfused for histology. The electrode was coated with dye (DiI, DiO, or DiD) for visualization, and recording locations were verified in most cases with histology post-hoc (**Extended Data Fig. 8a)**. In total, 19 recordings were made in 15 animals, with 132 MLIs recorded from the lateral cerebellum.

Data was recorded with SpikeGLX software (billkarsh.github.io) and potential units were identified using Kilosort 2.0^47^ and manually curated in phy (GitHub - cortex-lab/phy: phy: interactive visualization and manual spike sorting of large-scale ephys data). A custom plug-in to phy, phuyllum (from the Medina lab at Baylor College of Medicine) was used to support layer location identification for each contact along the probe. Well-isolated units have <5% refractory period violations in the ACG compared to the baseline firing rate and are missing <5% of spikes based on the unit amplitude histogram. Further analysis was carried out using custom Matlab programs. Voltage signals were filtered with a 300 Hz high pass (first order Butterworth) for waveform analysis. One hundred waveforms were extracted for waveform analysis, aligned to the trough, and the resulting average was normalized using a Euclidean norm.

#### Cell Identification

110 well-isolated neurons were positively identified as PCs based on the characteristic complex spike - simple spike pause (**Extended Data Fig. 8**). From these positively-identified PCs, we then identified units as MLIs in two ways: First, neurons that fired above 3 Hz and inhibited a complex spike-identified PC with a short latency (<4 ms) that were recorded on contacts in the PC or molecular layer were identified as MLIs. Second, neurons recorded on electrodes in the molecular layer, at least 40 μm from the PC layer, and that had a firing rate above 3 Hz were also identified as MLIs, regardless of any inhibition or lack thereof onto a PC (**Extended Data Fig, 8**). We impose this distance criteria for cells that do not inhibit PCs to avoid unintentionally categorizing any Purkinje cell layer interneuron as an MLI.

While PCs are known to provide inhibitory input to other PCs, we are confident we did not include any PCs in our MLI population. PC connections generally displayed enough synchrony to raise the baseline standard deviation such that the inhibition did not surpass 4*SD, which was our criteria for establishing an inhibitory connection (see below).

#### Rate-corrected cross correlograms for synchrony and inhibition characterization

Synchrony and inhibition were evaluated using rate-corrected cross correlograms^48^. Briefly, two neurons whose firing rates covary due to correlated inputs or state modulation will display lower frequency comodulation that can be visible on a ccg. We control for this possibility by constructing a “null ccg”, computed by assuming a uniform likelihood of spiking between any given pair of spikes in the spike train for one neuron in each pair. This null ccg shows how many spikes are expected at each time point given only the rate of each neuron. Subtracting the null ccg from the standard ccg gives us the rate corrected ccg, showing how many spikes/second are occurring above the expected coincident spikes given the time-varying firing rates of each neuron. This rate-corrected ccg shows the true amount of synchrony (or inhibition) between neurons, irrespective of low frequency comodulation of firing rates. All correlograms were computed using only time periods during the recording when the animal was quiescent (neither moving nor licking unless otherwise noted). Neurons were classified as synchronous when their rate-corrected ccg surpassed 4 SD above baseline at time zero, and as inhibitory when their rate-corrected ccg dipped below 4 SD from baseline between 0 and 4 ms. (For synchrony evaluations baseline was calculated from -20 to -5 ms on the ccg, and for inhibitory evaluations baseline was calculated from -20 to 0 ms on the ccg.)

### Statistics and reproducibility

We did not use statistical methods to pre-determine sample sizes. Technical limitations made it only feasible to analyze one mouse for EM analysis in this study. Details of statistical tests for **Fig. 1** and **Fig. 2** are summarized in **Table 1**. Statistical significance was assumed at *p*<0.05, and exact *p* and *n* values are stated in the figure legends and **Table 1**. For *in vivo* recordings, population firing statistics were compared with violin plots constructed in Matlab (Hoffmann H, 2015: violin.m - Simple violin plot using Matlab default kernel density estimation. INRES (University of Bonn), Katzenburgweg 5, 53115 Germany.). Kernel densities on the same panel were estimated with common bandwidth supported from the minimum-5 to the maximum+5 of the grouped data (assuming normal density).

**Table 1.**
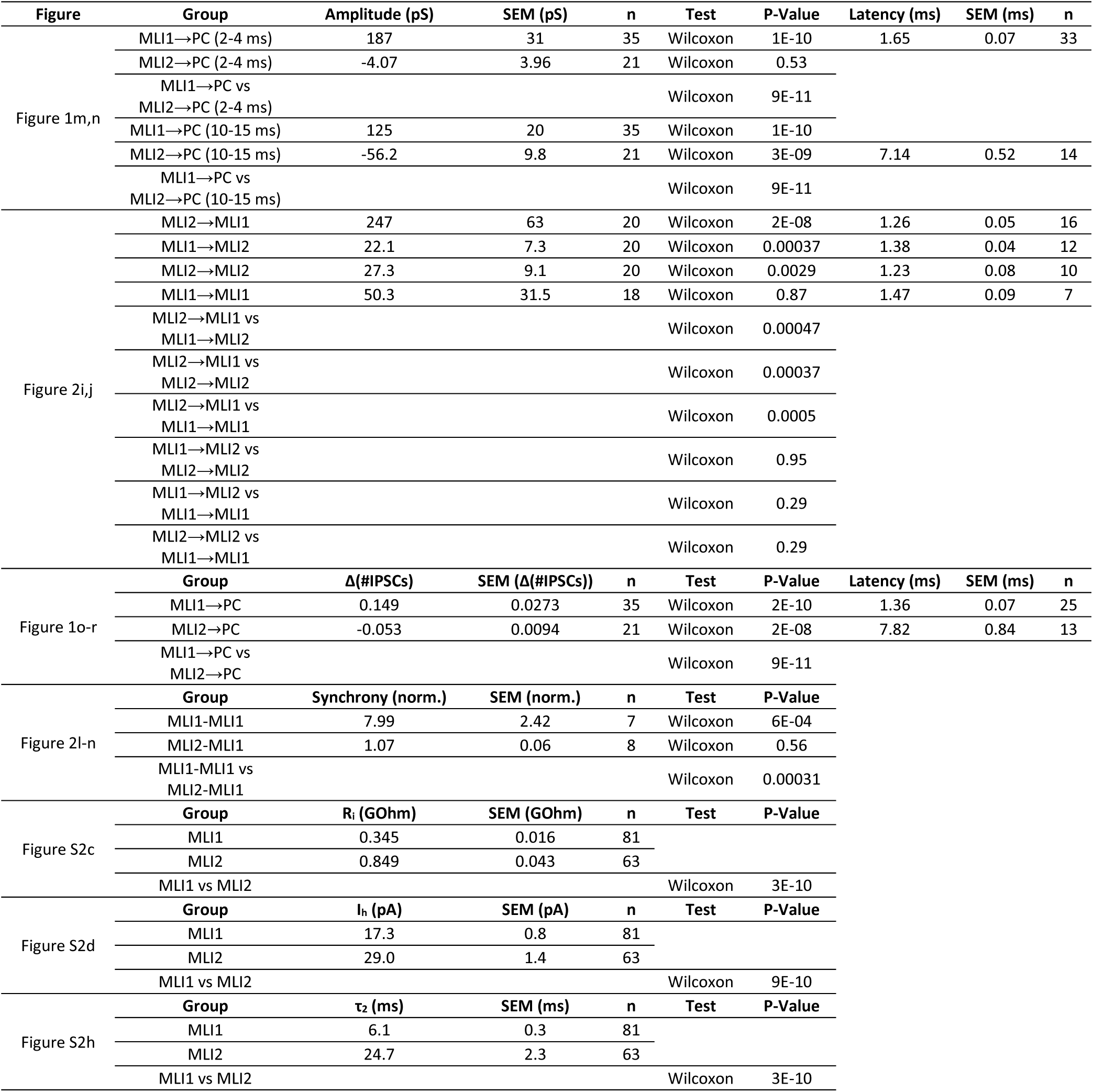
Summary and statistics of slice electrophysiology experiments.

Statistical comparisons of firing rates between cell type populations were performed with a two-sample Kolmogorov-Smirnov test (Matlab). Comparisons of differences in firing rates between states were computed using the Wilcoxon rank sum test (Matlab). Data are reported as mean ± standard error.

## Data availability

The electron microscopy dataset is publicly available at the BossDB (https://bossdb.org/) repository https://bossdb.org/project/nguyen_thomas2022. Further directions for accessing neuron 3D renderings and derived neuron connectivity graphs are available at https://github.com/htem/cb2_project_analysis. Source data will be provided.

## Code availability

Software used in this work is open-source and available in the following repositories. Daisy: https://github.com/funkelab/daisy. MD-Seg front-end: https://github.com/htem/neuroglancer_mdseg/ tree/segway_pr_v2. MD-Seg back-end: https://github.com/htem/segway.mdseg. Segmentation and synapse prediction scripts: https://github.com/htem/segway. Analysis code: https://github.com/htem/cb2_project_analysis. SpikeGLX software: https://billkarsh.github.io/SpikeGLX. Kilosort 2.0: https://github.com/cortex-lab/Kilosort. phy: https://github.com/cortex-lab/phy.

## Acknowledgements

We thank Shuting Wu, Joon-Hyuk Lee and Vincent Huson for comments on the manuscript. We thank David Herzfeld for analysis assistance with rate corrected CCGs, and Javier F. Medina, Francisco Naveros and Alvaro Sanchez-Lopez for the use of the Phy plugin "Phyllum" to corroborate layer identification for neuropixels recordings. This work was supported by the NIH (R01MH122570 and R35NS097284 to W.G.R., F32NS133036 to E.P.L., R21NS085320 and RF1MH114047 to W.-C.A.L., 1U19MH114821 to E.Z.M., R01NS128054 and R01NS112917 to C.A.H.), the Bertarelli Program in Translational Neuroscience and Neuroengineering, Stanley and Theodora Feldberg Fund, and the Edward R. and Anne G. Lefler Center. Portions of this research were conducted on the O2 High Performance Compute Cluster at Harvard Medical School partially provided through NIH NCRR (1S10RR028832-01) and a Foundry Award for the HMS Connectomics Core. Equipment in the HMS Neurobiology Imaging Facility was used for confocal imaging. We thank Brad Lowell, Daqing Wang and the BNORC transgenic core for help in making *Nxph1^Cre^* mice.

## Author contributions

W.G.R., L.M. and E.P.L. conceptualized the project and designed the slice experiments. W.G.R. and W.-C.A.L. conceptualized and designed EM experiments. C.A.H. conceptualized and designed *in vivo* experiments. E.Z.M. provided crucial insights into MLI1s and MLI2s in the initial phase of this project. E.P.L. and L.M. performed slice experiments and analyzed slice data. L.M. made *Nxph1^Cre^* mice using the BNORC transgenic core. E.P.L. performed smFISH experiments and analyzed smFISH data. T.N. generated the automated segmentations in the serial EM dataset and helped guide EM reconstructions. W.-C.A.L. provided resources and infrastructure for EM reconstructions. A.N. performed EM reconstructions and analyzed the EM data. T.O. used fluorescent fills of MLIs to determine the characteristic morphological features of MLI subtypes that were crucial to identifying them in EM reconstructions. M.H. conducted and analyzed *in vivo* experiments. W.G.R. and E.P.L. wrote the paper with input from the other authors. C.A.H. and M.H. wrote the initial draft of the text associated with *in vivo* experiments.

