## Extended Data Figures for "Cerebellar circuits for disinhibition and synchronous inhibition"

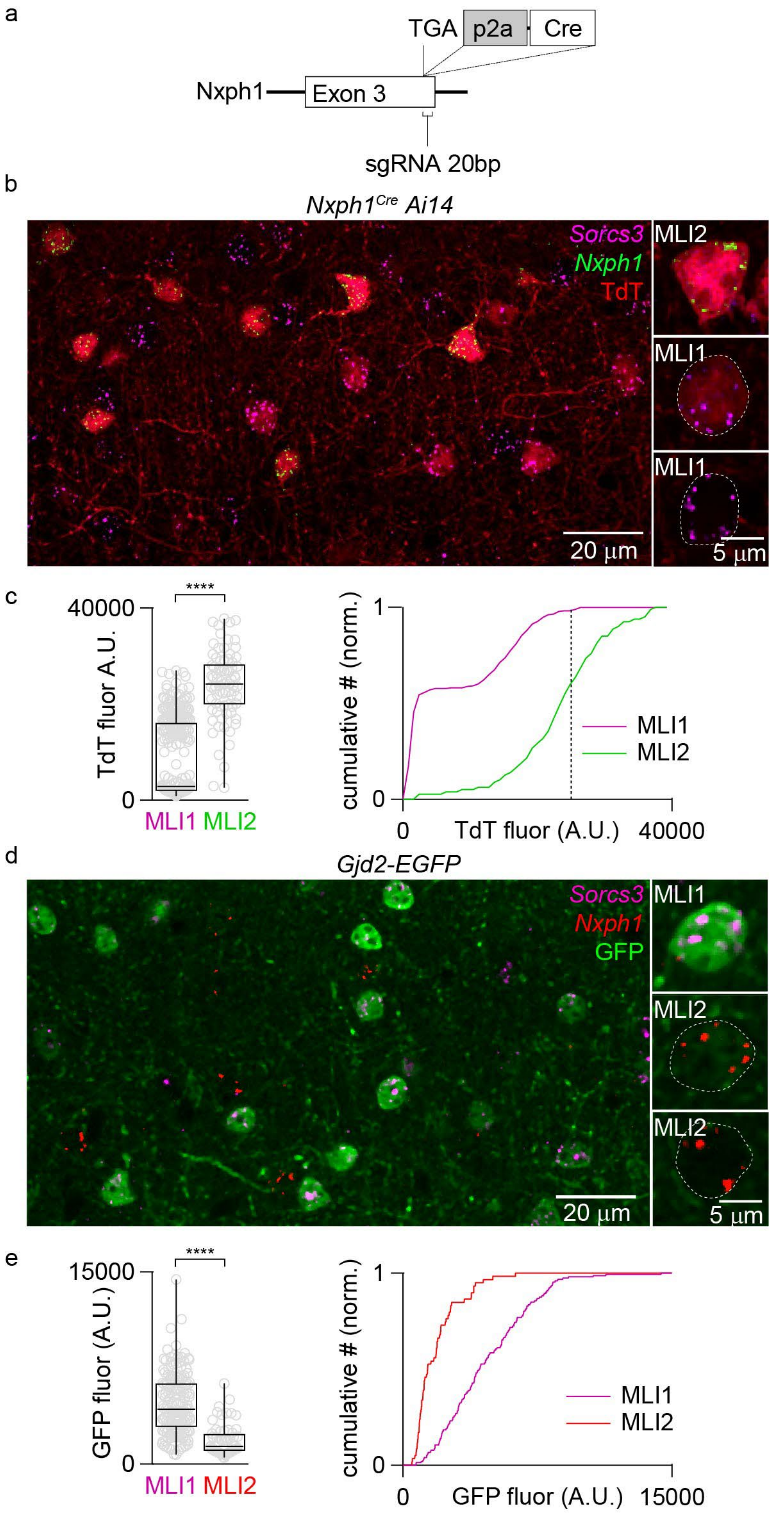


**Extended Data Fig. 1. Characterization of *Nxph1$^{Cre}$* mice and *Gjd2-EGFP* mice**

a. Schematic summarizing the strategy used to make an *Nxph1$^{Cre}$* mouse.

b. Fluorescence image of a region of the cerebellar molecular layer of an *Nxph1$^{Cre}$ Ai14* mouse. *In situs* were performed to identify MLI1s (Sorcs3+, *purple*) and MLI2s (Nxph1+, *green*). Expanded views of three cells are shown.

c. (*left*) The TdT fluorescence of the somata are shown for Sorcs3+ cells (MLI1, n=289) and Nxph1+ cells (MLI2, n=79) ($p$=1E-09). (*right*) Normalized cumulative plot of the TdT fluorescence data in c. Dashed line is used as a threshold showing that most bright cells are MLI2s: 0.13% of MLI1s and 36.6 % of MLI2s had a fluorescence intensity of > 26000 A.U.s.

d. Fluorescence image of a region of the cerebellar molecular layer of a *Gjd2-EGFP* mouse. *In situs* were performed to identify MLI1s (Sorcs3+, *purple*) and MLI2s (Nxph1+, *red*). Expanded views of three cells are shown.

e. (*left*) The GFP fluorescence intensities of the somata are shown for Nxph1+ cells (MLI2, n=59) and Sorcs3+ cells (MLI1, n=152) ($p$=1E-09).

(*right*) Normalized cumulative plot of the GFP fluorescence data in e.

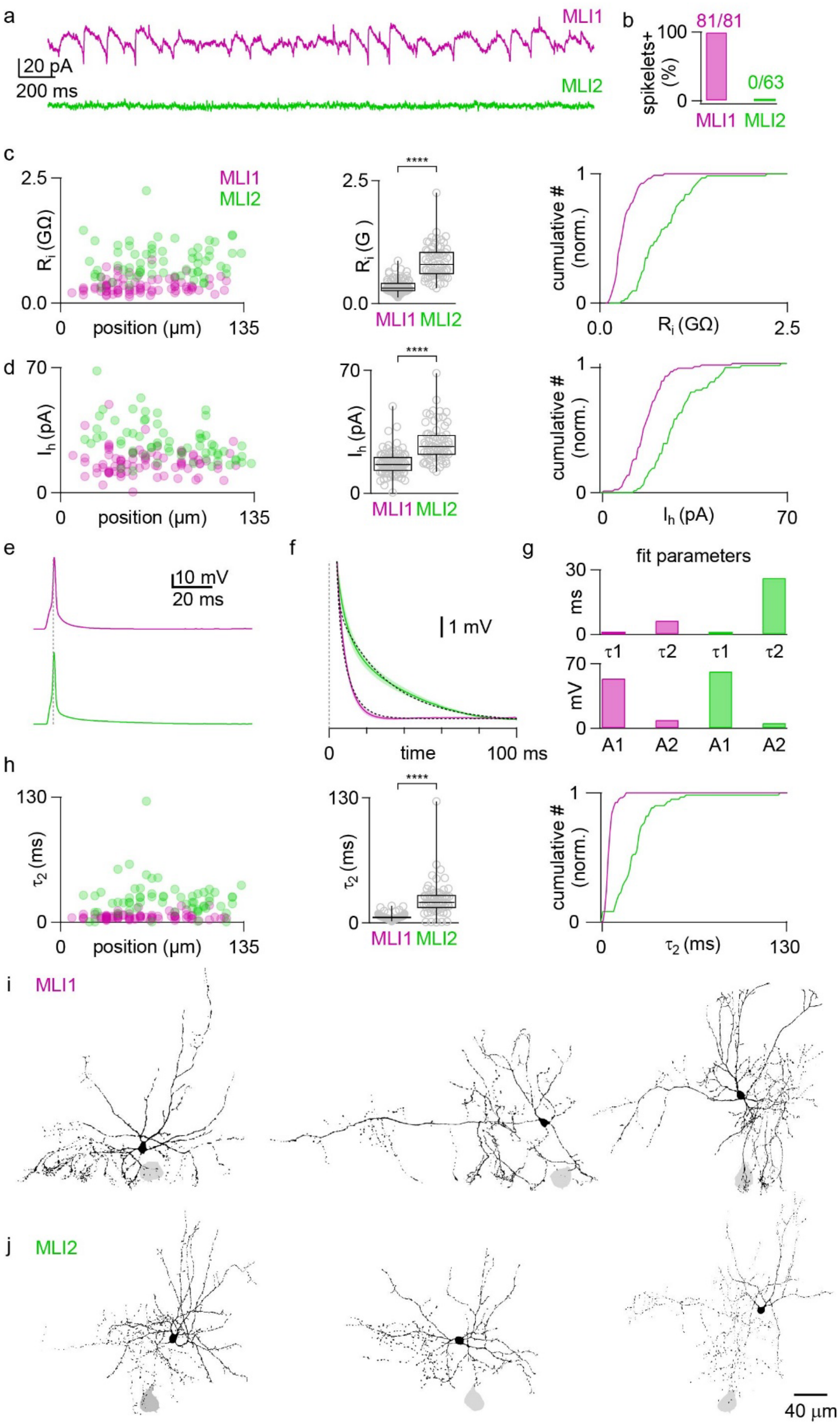


**Extended Data Fig. 2. Approach taken to identify MLI1s and MLI2s.**

a. The presence of spikelets was used to identify MLI1s.

b. Spikelets were present in all MLI1s and absent in all MLI2s included in Fig. 1 and Fig. 2. Additional criteria were used to exclude MLI1s that lacked spikelets (Kozareva et al 2021).

c. MLI2s tended to have a higher resistance than MLI1s (*p*=3E-10 ).

d. MLI2s tended to have a larger $I_h$ than MLI1s (*p*=9E-10), particularly for cells near the PC layer.

e. Example traces for action potentials evoked in an MLI1 and an MLI2.

f. Average traces as in e for all identified MLI1s and MLI2s.

g. Membrane potential was fit with a double exponential decay for the traces in f, and the fit parameters are summarized.

h. Similar double exponential fits were performed for all MLIs, and the slow time constant is shown for each cell as a function of position. The slow time constant was significantly slower in MLI2s (*p*=3E-10), as shown in the box plot and the cumulative plot.

i. Fluorescent labelling of MLI1s from paired MLI1-PC recordings, with the only the PC somata shown.

j. As in **i**, but for MLI2s.

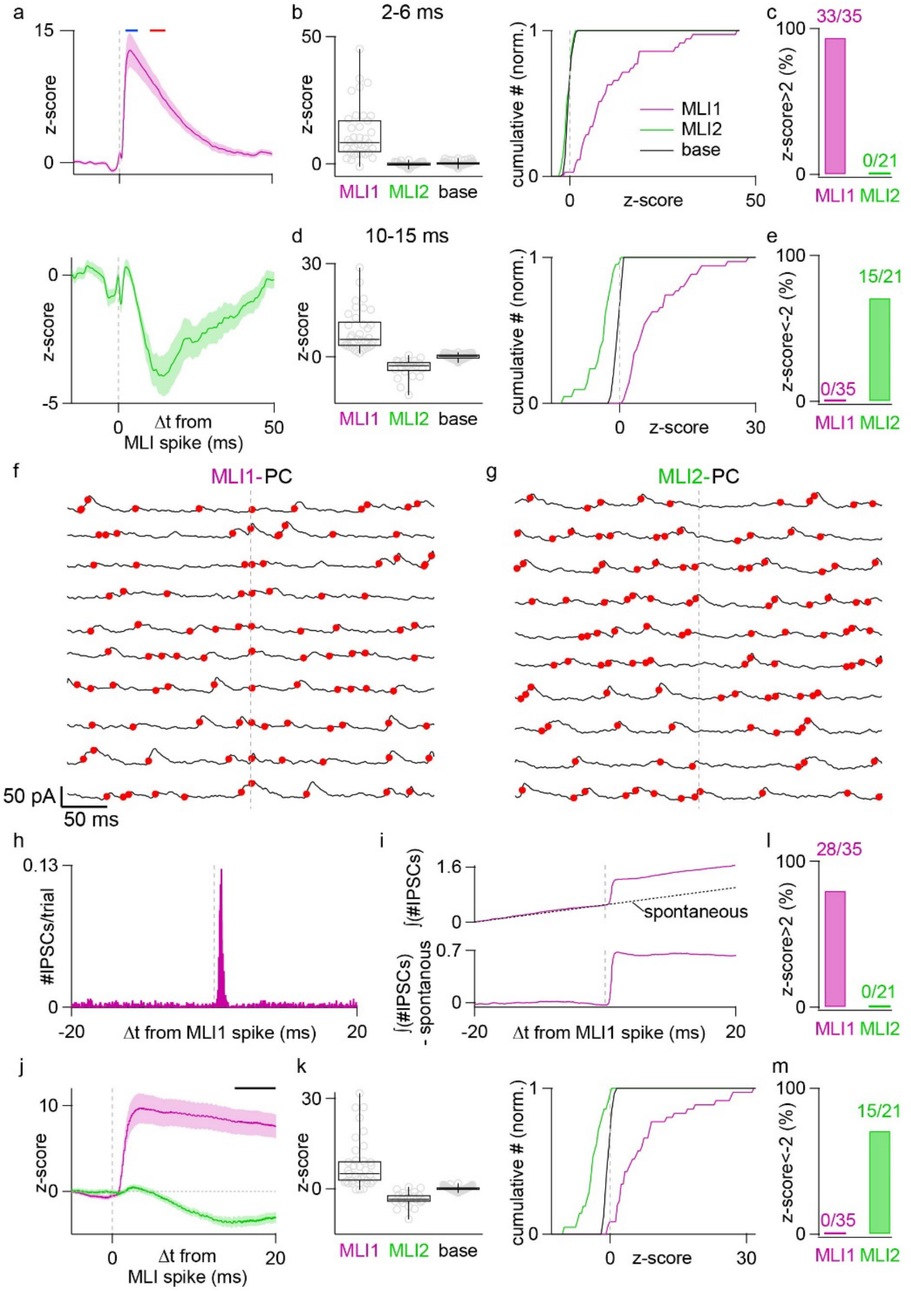


**Extended Data Fig.3. Analysis of evoked and spontaneous IPSCs in Purkinje cells.**

a. Average z scores for MLI1-PC pairs and MLI2-PC pairs. The blue bar (2-6 ms) and red bar (10-15 ms) show the regions used for analysis.

b. Box plot and cumulative summary for 2-6 ms interval following the presynaptic action potential. Base refers to a similar analysis of the baseline in the absence of stimulation.

c. The z-score (>2) was used to determine the fraction of MLI-PC with a monosynaptic connection.

d. As in b, but for 10-15 ms following stimulation.

e. The z-score (<-2) was used to determine the fraction of MLI-PC pairs where disinhibition was observed.

f. Traces recorded from a PC during a paired MLI1-PC recording. Traces are aligned to the time of an action potential evoked in the presynaptic MLI (vertical dashed line). The detected IPSCs are indicated by solid red symbols. These consecutive traces are 10 of the 500 traces used to determine the raster plot in Fig. 1f.

g. Same as f, but for an MLI2-PC pair. These consecutive traces are 10 of the 500 hundred traces used to determine the raster plot in Fig. 1l.

h. Histogram showing the number of IPSCs detected per trial per 50 µs bin for an MLI1-PC pair.

i. (*top*) Integration of the histogram in h. The dashed line is a linear fit to the -200 to 0 ms interval that reflects the spontaneous events. (*bottom*) A plot of the integrated events – the dashed line (spontaneous events) was used to determine the effects of stimulation of the frequency of IPSCs.

j. The average z-score for the integrated events due to stimulation is shown for all MLI-PC pairs. A bar shows the interval (15-20 ms) used for analysis in k.

k. Summary of the z-scores for all MLI-PC pairs for the effect of stimulation on the integrated number of IPSCs per spike, with a box plot shown on the left and a cumulative plot shown on the right.

l. Summary of the MLI-PC pairs from k that had a z-score >2, consistent with monosynaptic inhibition.

m. Summary of the MLI-PC pairs from k that had a z-score <-2, consistent with disinhibition.

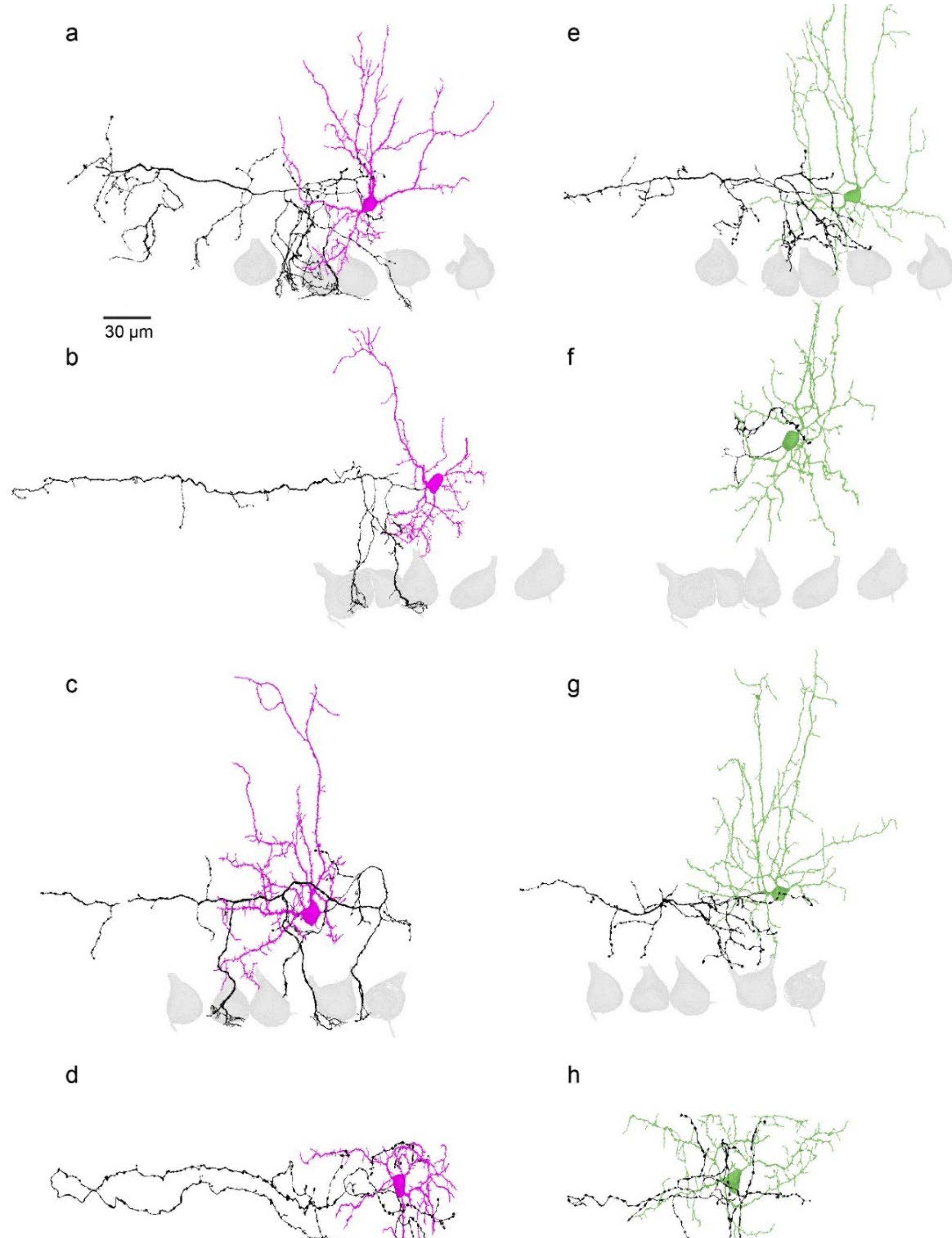


**Extended Data Fig. 4. Examples of EM reconstructions of MLIs.**

**a-d).** MLI1s

**e-h).** MLI2s

Four MLI1s and four ML2s were matched to allow the comparison of morphologies of pairs of cells located at approximately the same distance away from the PC layer (**a, e; b, f; c, g; d, h**). MLIs in **d** and **h** are located near the top of the molecular layer.

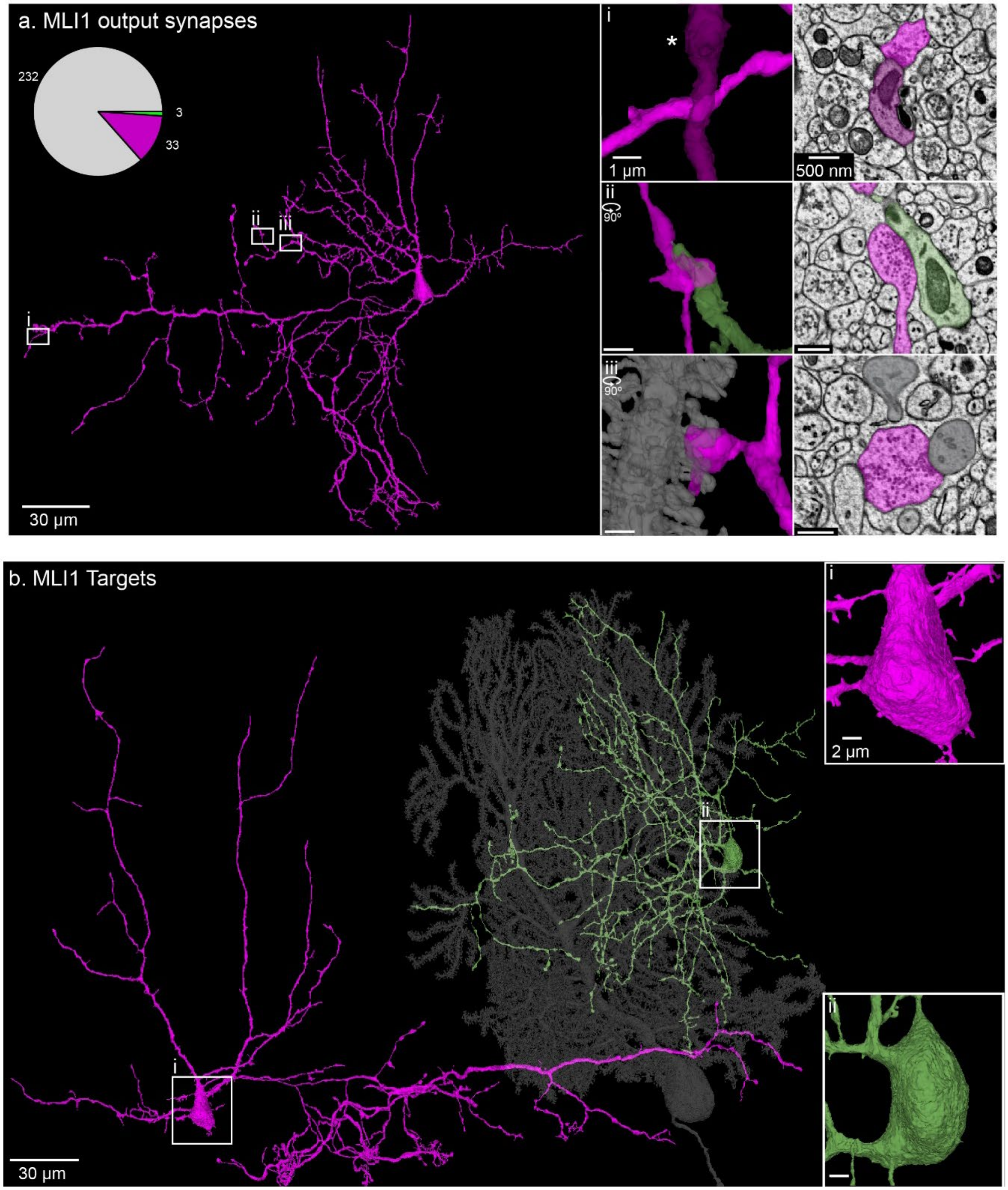


**Extended Data Fig. 5. Examples of reconstructions of MLI1 synaptic connections onto different targets.**

**a.** An image of a reconstructed MLI1. The pie chart summarizes the synaptic connections made onto different types of targets. The 3 regions (**i-iii**) are shown on expanded scales with both reconstructed views and an EM through the synaptic contact. Examples of an MLI1-MLI1 synapse (i), an MLI1-MLI2 synapse (ii) and an MLI1-PC synapse (iii) are shown.

**b.** Reconstructions of three of the neurons targeted by this MLI1 are shown: MLI1 (*purple*), MLI2 (*green*) and PC (*grey*). Expanded views of the somata of the MLI targets show the smooth cell body characteristic of an MLI2 (*green*) and the spiny cell body characteristic of an MLI1 (*purple*).

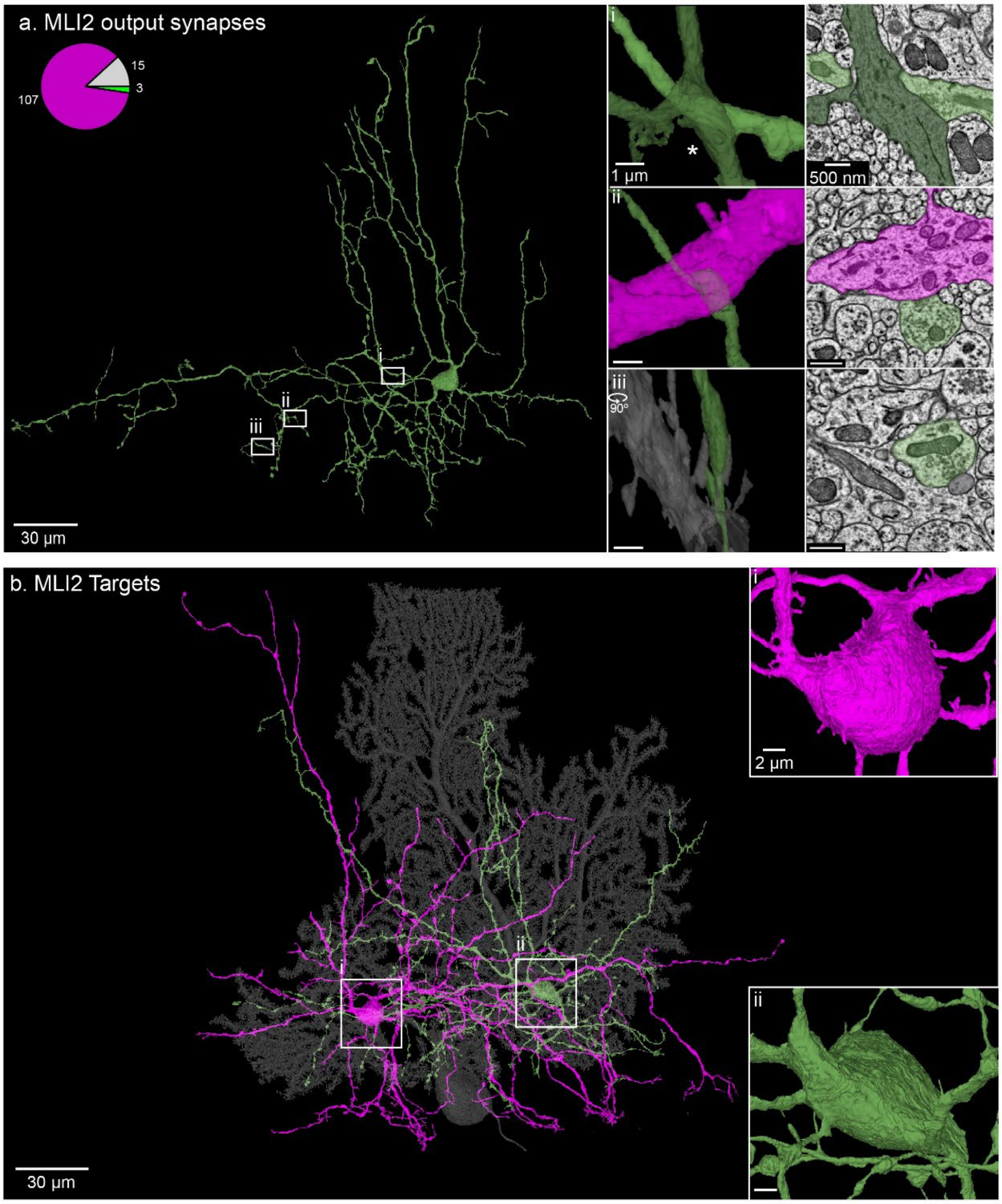


**Extended Data Fig. 6. Examples of reconstructions of MLI2 synaptic connections onto different targets.**

- **a.** An image of a reconstructed MLI2. The pie chart summarizes the synaptic connections made onto different types of targets. The 3 regions (**i-iii**) are shown on expanded scales with both reconstructed views and an EM through the synaptic contact. Examples of an MLI2-MLI2 synapse (i), an MLI2-MLI1 synapse (ii) and an MLI2-PC synapse (iii) are shown.
- **b.** Reconstructions of three of the neurons targeted by this MLI2 are shown: MLI1 (*purple*), MLI2 (*green*) and PC (*grey*). Expanded views of the somata of the MLI targets show the smooth cell body that is characteristic of an MLI2 (*green*), and the spiny cell body that is characteristic of an MLI1 (*purple*).

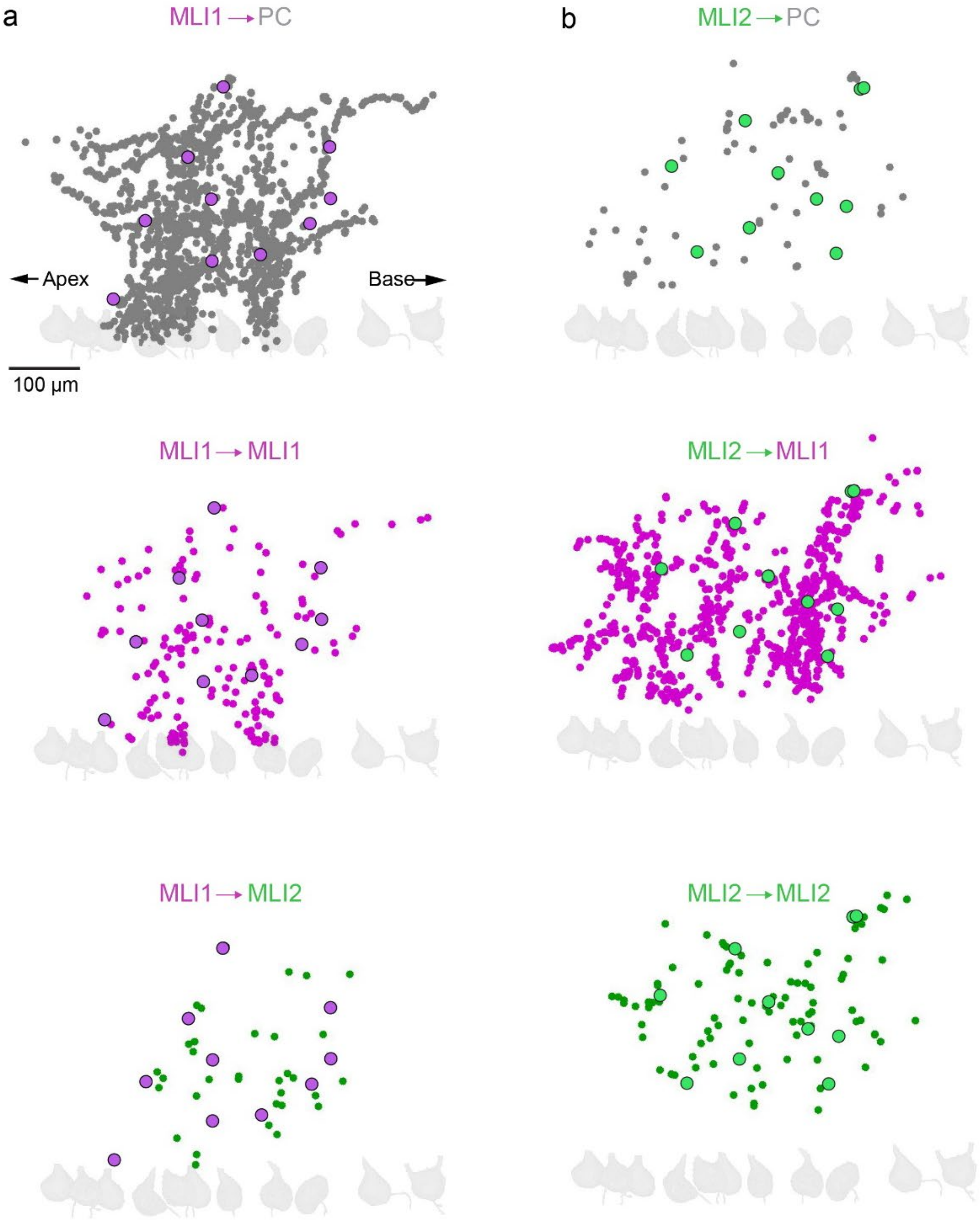


**Extended Data Fig. 7. Position of MLI1 and MLI2 synapses.**

- **a.** The positions of synapses made by MLI1s onto different types of targets are indicated for the 10 cells shown in **Fig. 4a**. The positions of the cell bodies are indicated (*large purple circles*). These data are replotted in **Fig. 4c**, but with the cell bodies superimposed and the relative positions of their synapses adjusted accordingly.
- **b.** The positions of synapses made by MLI2s onto different types of targets are indicated for the 10 cells shown in **Fig. 4b**. The positions of the cell bodies are indicated (*large green circles*). These data are replotted in **Fig. 4e**, but with the cell bodies superimposed and the relative positions of their synapses adjusted accordingly.

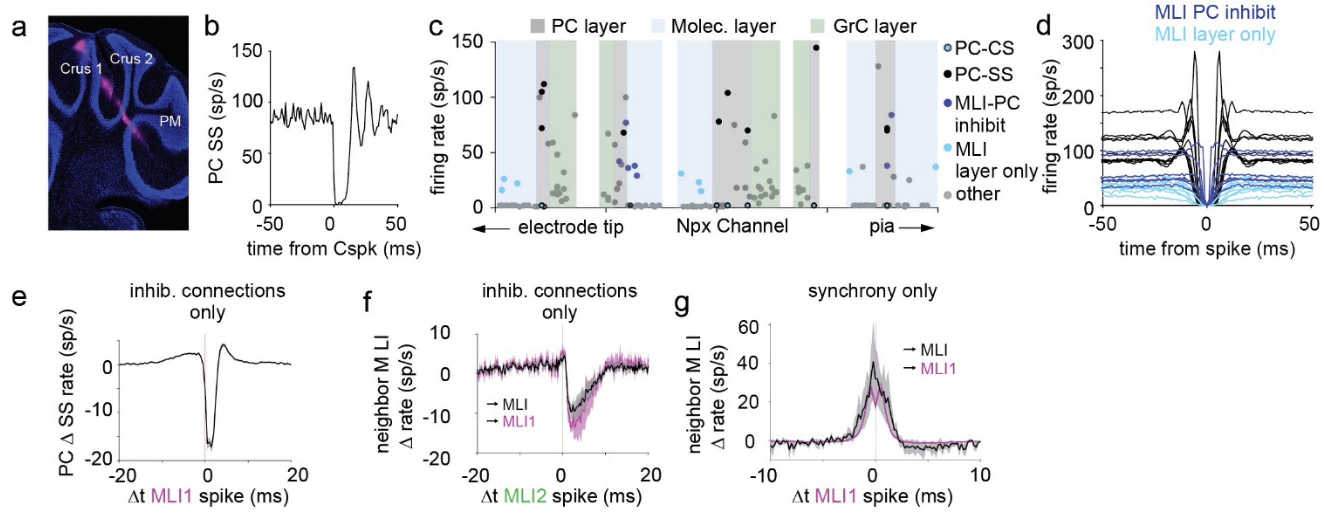


**Extended Data Fig. 8. *In vivo* PC and MLI identification and properties.**

**a.** Labeled Neuropixels tract for recording through several layers of the cerebellum.

**b.** The complex spike pause of an identified PC.

**c.** Firing rates of all well-isolated units from the recording in panel a. demonstrate a clearly identifiable laminar structure. Pause-identified PCs are highlighted in black. Laminar structure based on PC identification and surrounding unit firing rates. The molecular layers (blue background) are characterized by many 1 Hz “fat spikes” that are the complex spike recordings from the PC dendrites. Other units isolated in these layers (>3 Hz) are identified as MLIs. PC layers are identifiable by numerous high-firing rate units, and the presence of units with simple spikes and complex spikes.  The granular layer contains neither fat spikes nor simple spikes but contains units that fire at a range of intermediate frequencies. MLIs were identified by CCGs with a significant depression in a pause-identified PC (blue), or by their presence in the molecular layer away from any potential PC (cyan).

**d.** Auto-correlograms of pause-identified PCs and MLIs from the recording in **a** and **c**.

**e.** Mean inhibition of MLI1-PC inhibitory pairs (203 pairs). By definition, there are no MLI2-PC inhibitory pairs.

**f.** MLI2-MLI inhibitory pairs (9 total MLI2-MLI pairs, 5 MLI2-MLI1 pairs, 0 MLI2-MLI2 pairs).

**g.** Mean cross-correlograms of MLI1-MLI synchronous pairs (125 MLI1-MLI1 pairs, 5 MLI1-unknown MLI pairs).
